# Oxidoreductases generate hydrogen peroxide that drives iron-dependent lipid peroxidation during ferroptosis

**DOI:** 10.1101/2020.08.01.231993

**Authors:** Bo Yan, Youwei Ai, Ze Zhang, Qi Sun, Yan Ma, Zhiyuan Zhang, Xiaodong Wang

## Abstract

The inhibition of antioxidant systems of glutathione peroxidase 4 (GPX4) or ferroptosis suppressor protein 1 (FSP1) causes iron-dependent peroxidation of polyunsaturated phospholipids that leads to cell death, a process known as ferroptosis. The mechanisms underlying iron-dependent lipid peroxidation are under active debate. Here, we report that two endoplasmic reticulum-residing oxidoreductases, NADPH-cytochrome P450 reductase (POR) and NADH-cytochrome b5 reductase (CYB5R1), are responsible for the iron-dependent peroxidation of polyunsaturated phospholipids and membrane disruption that executes ferroptosis. Genetic ablation of POR and CYB5R1 or mutations that eliminate POR’s electron transfer activity blocked ferroptosis. *In vitro* enzymatic assays established that POR and CYB5R1 catalyze hydrogen peroxide production by transferring electrons from NADPH/NADH to oxygen, which is then used to carry out iron-dependent lipid peroxidation via a Fenton reaction. The lipid peroxidation reaction catalyzed by POR and CYB5R1 additively disrupts polyunsaturated phospholipid-containing liposomes. Finally, POR knockdown confers significant protective effects during concanavalin A-induced, ferroptosis-associated acute liver injury *in vivo.* Our study thus indicates that POR and CYB5R1 are the enzymes of the “oxidant” system that operates to contravene the antioxidant GPX4/FSP1 systems; the balance between these two systems determines cell commitment to ferroptosis.

## INTRODUCTION

Ferroptosis is a form of regulated necrotic cell death that has been implicated in a variety of medical disorders including neurodegenerative diseases, acute tissue failure, and cancer suppression^1, 2^. Ferroptosis is characterized by the iron-dependent accumulation of lipid peroxides on cell membranes^1, 3^. Mechanistically, several proteins have been shown to regulate ferroptosis by limiting the production of lipid peroxides: cystine-glutamate antiporter (xCT)^1^, Glutathione peroxidase 4 (GPX4)^3^, ferroptosis suppressor protein 1 (FSP1)^4, 5^, and GTP cyclohydrolase-1 (GCH1)^6^. xCT is a cystine antiporter that provides cells with cysteine to synthesize glutathione, an antioxidant and substrate for GPX4. GPX4 glutathione-dependently catalyzes the reduction of lipid peroxides, thereby actively preventing ferroptosis. Direct inhibition of GPX4 or depletion of its substrate glutathione can trigger ferroptosis. FSP1 and GCH1 catalyze lipid peroxide reduction in a glutathione-independent manner by reducing Coenzyme Q10 to ubiquinol and synthesizing 6(R)-L-erythro-5,6,7,8-tetrahydrobiopterin respectively, which can then trap lipid peroxides.

When GPX4 or FSP1 is defective, lipid peroxides accumulate in cells, reaching lethal levels that result in ferroptosis^7^. Given that polyunsaturated-fatty-acid-containing phospholipids (PUFA phospholipids) are known to be substrates for lipid peroxidation, reducing cellular PUFA phospholipid content by knocking down PUFA phospholipid synthesis enzymes such as acyl-CoA synthetase long-chain family member-4 (ACSL4) inhibits ferroptosis^8^. AMPK-ACC and NF2-YAP signaling pathways are also known to impact ferroptosis by regulating PUFA metabolism and cellular phospholipid composition^9, 10^. However, the enzymes specifically involved in the lipid peroxidation process remain unclear. Previous studies have suggested that lipoxygenases (ALOXs)—particularly 15ALOX—drive lipid peroxidation for ferroptosis^11, 12^. Recently, however, the necessity for ALOX involvement in ferroptosis was called into question by the discovery of previously unknown radical-trapping activities for ALOX-inhibiting small molecules^13^. Thus, there must be other components that catalyze lipid peroxidation during ferroptosis.

Here, we used a genome-wide, unbiased CRISPR-Cas9 screen alongside a small-scale targeted reductase knockdown screen and found that NADPH-cytochrome P450 reductase (POR) and NADH-cytochrome b5 reductase 1 (CYB5R1) directly catalyze lipid peroxidation and contribute to ferroptosis execution. Intriguingly, the lipid peroxidation reactions catalyzed by these two enzymes represent an oxygen-dependent hijacking of their normal oxidative functions to produce hydrogen peroxide, which can then produce hydroxyl radicals to drive lipid peroxidation. Based on liposome leakage experiments, we confirm that POR- and CYB5R1-catalyzed lipid peroxidation directly disrupts the integrity of PUFA-containing phospholipid membranes. Finally, we found that POR knockdown in mouse liver confers protective effects during acute liver injury caused from ferroptosis, indicating that inhibition of POR represents a new therapeutic target for the treatment of ferroptosis-related diseases.

## RESULTS

### PACMA31 induces ferroptosis by targeting GPX4

During our ongoing study of protein disulfide isomerase (PDI) inhibitors, we serendipitously found that an irreversible PDI inhibitor, PACMA31^14^, induced cell death that could be fully blocked by ferroptosis inhibitor Fer-1 but not by apoptosis or necroptosis inhibitors (Fig.1a and Extended Data Fig.1a), indicating PACMA31 was a ferroptosis trigger. PACMA31-induced ferroptosis was independent of its inhibition of PDI, as genetic knockdown of PDI did not cause cell death (Extended Data Fig.1b-c). We subsequently equipped PACMA31 with an alkyne handle on an ester group (“PACMA31-Probe”, Extended Data Fig.1d) and after confirming that PACMA31-Probe still induced ferroptosis (Extended Data Fig.1e), we used the probe to pull down cellular interactive proteins of PACMA31-Probe and identified GPX4 as one of its targeting proteins (Fig.1b). Both PACMA31 and the previously known irreversible GPX4 inhibitor RSL3, which binds to the selenocysteine of GPX4^11^, could outcompete PACMA31-Probe for binding with GPX4 (Fig.1b). An LC-MS/MS analysis of PACMA31 reaction products further confirmed the covalent binding of PACMA31 to the active site cysteine of a GPX4 variant (Extended Data Fig.1f), suggesting that ferroptosis caused by PACMA31 is due to its direct targeting and inhibition of GPX4.

**Fig. 1.**
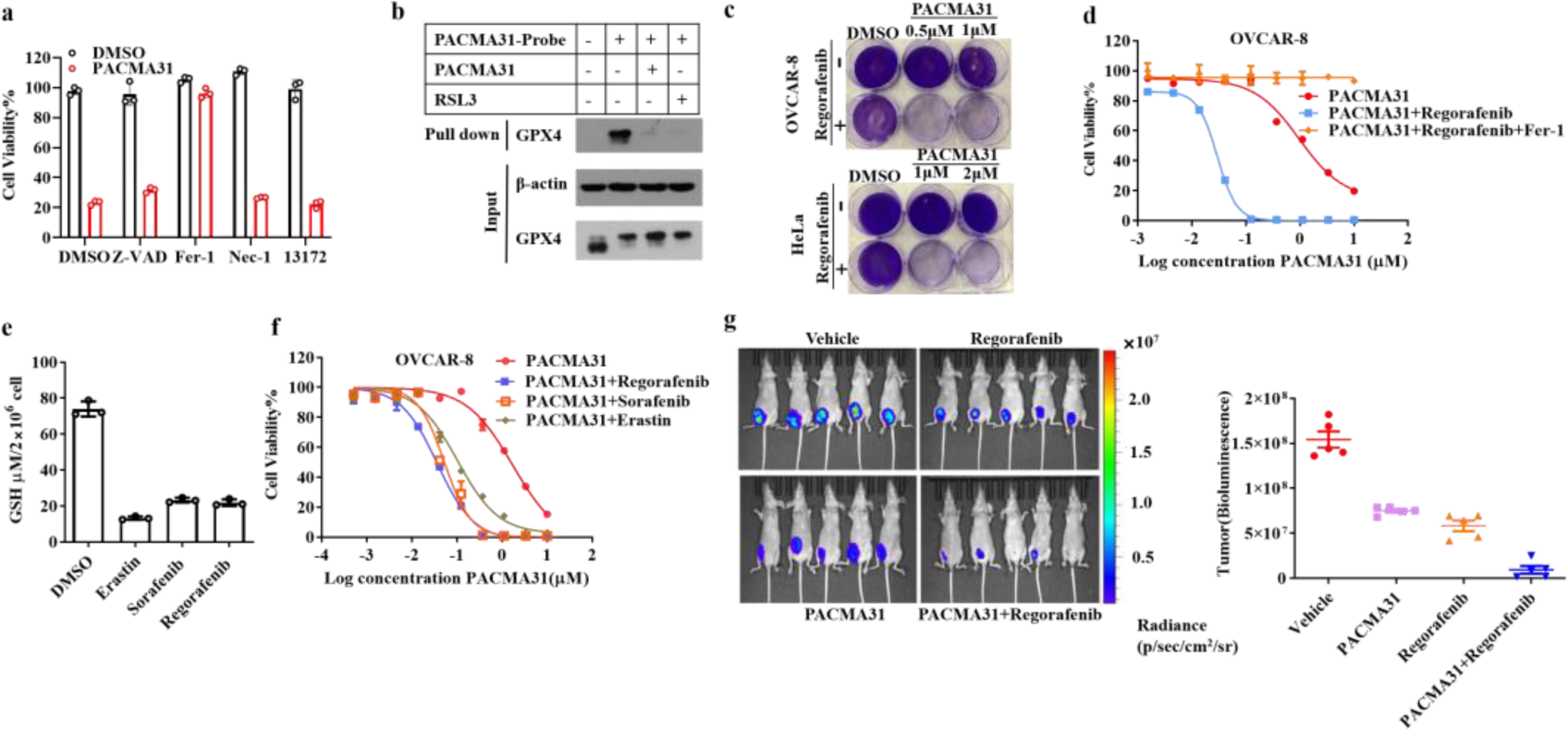
Regorafenib synergizes with PACMA31 to induce ferroptosis. (a) The cell viability of OVCAR-8 cells treated with PACMA31 (10 μM) alone or combined with the apoptosis inhibitor Z-VAD-fmk (10 μM), the ferroptosis inhibitor Fer-1 (1 μM), or necroptosis inhibitors, including the receptor interacting protein kinase 1 (RIP1) inhibitor Nec-1 (1 μM) and the mixed lineage kinase domain-like protein (MLKL) inhibitor 13172 (200 nM). (b) PACMA31 binds GPX4. Protein levels were analyzed by western blotting using antibodies against GPX4 and β-actin. (c) Methylene blue staining showing the colony-forming of OVCAR-8 and HeLa cells with indicated treatments. (d) The cell viability curves for OVCAR-8 cells treated with different doses of PACMA31 alone or combined with regorafenib (10 μM) or combined with regorafenib and Fer-1 (1 μM). (e) The glutathione levels in OVCAR-8 cells with indicated treatments for 24h (Erastin 1 μM; regorafenib 10 μM; sorafenib 10 μM). (f) The cell viability curves for OVCAR-8 cells treated with different doses of PACMA31 alone or combined with Erastin (1 μM), regorafenib (10 μM) or sorafenib (10 μM). (g) Bioluminescence imaging of mice bearing luciferase-expressing OVCAR-8 xenografts, performed after administration with the indicated drugs for 23 days. Quantitative analysis of the bioluminescence in OVCAR-8 xenografts mice is presented as total photon flux according (assessed using IVIS software). n = 5 mice each group. In this figure, cell viability assays were examined 12h after treatment by measuring the ATP level (ATP level of DMSO samples set to 100%). Bar graphs show the mean ± SD. Student’s t test (two-tailed, unpaired) was used for the comparison of the indicated two groups: *p < 0.05; **p < 0.01; ***p< 0.001; ns, not significant.

### Regorafenib synergizes with PACMA31 in ferroptosis

We performed a systematic library screen comprising of 272 FDA-approved and presently-investigated anti-cancer drugs for compounds that would work synergistically with PACMA31 for ferroptosis induction in OVCAR-8 cells and identified regorafenib as the most potent compound (Extended Data Fig.2a). Additional experiments with several cancer cell lines indicated that the synergistic effect of PACMA31 with regorafenib is general (Fig.1c and Extended Data Fig.2b-c). Similar to PACMA31, RSL3 also showed synergistic effects with regorafenib (Extended Data Fig.2d). Cell death induced by the co-treatment of PACMA31 and regorafenib (referred to subsequently as P+R) was fully blocked by Fer-1 (Fig.1d).

Since regorafenib is structurally similar to sorafenib^15^, an inhibitor of xCT that can drive ferroptosis by causing depletion of cellular glutathione (GSH)^16^, it was not surprising that regorafenib significantly depleted GSH levels in cells (Fig.1e). Other xCT inhibitors such as sorafenib and Erastin also exerted synergistic effects with PACMA31 to kill OVCAR-8 cells at sub-toxic concentrations (Fig.1f). These observations suggested that the synergistic effect of regorafenib with PACMA31 was due to xCT inhibition. The P+R co-treatment also resulted in a marked reduction in the growth of OVCAR-8 tumor xenografts *in vivo* as measured by bioluminescence imaging (Fig.1g and Extended Data Fig2.e-g). The combination of P+R thus provided us a potent ferroptosis inducer both *in vitro* and *in vivo*.

### POR is required for ferroptosis

Based on the robust ferroptosis trigger (P+R), we performed unbiased genome-wide CRISPR-Cas9 screens in HeLa and OVCAR-8 cells treated with lethal P+R concentration (Fig.2a). Our results indicated that *ACSL4*, a known ferroptosis regulator involved in polyunsaturated fatty acid (PUFA)-phospholipids synthesis^8^, was among the most highly enriched candidate genes, together with five other candidates (*POR*, *ATL3*, *DTYMK*, *HHAT*, and *INTS6*) (Extended Data Fig.3a). We knocked down the expression of these genes using 2-3 different shRNAs and found that only the blockage effect of knocking down ACSL4 and POR was reproducible (Extended Data Fig.3b-m). Additionally, knockdown of POR in HeLa, HT-1080, and murine MEF cells all resulted in resistance to P+R induced ferroptosis (Extended Data Fig.4a-f). Furthermore, the role of POR in P+R induced ferroptosis could be validated by its knockout with additional sgRNAs of different sequences (Fig.1b-c and Extended Data Fig.4g-h).

**Fig. 2.**
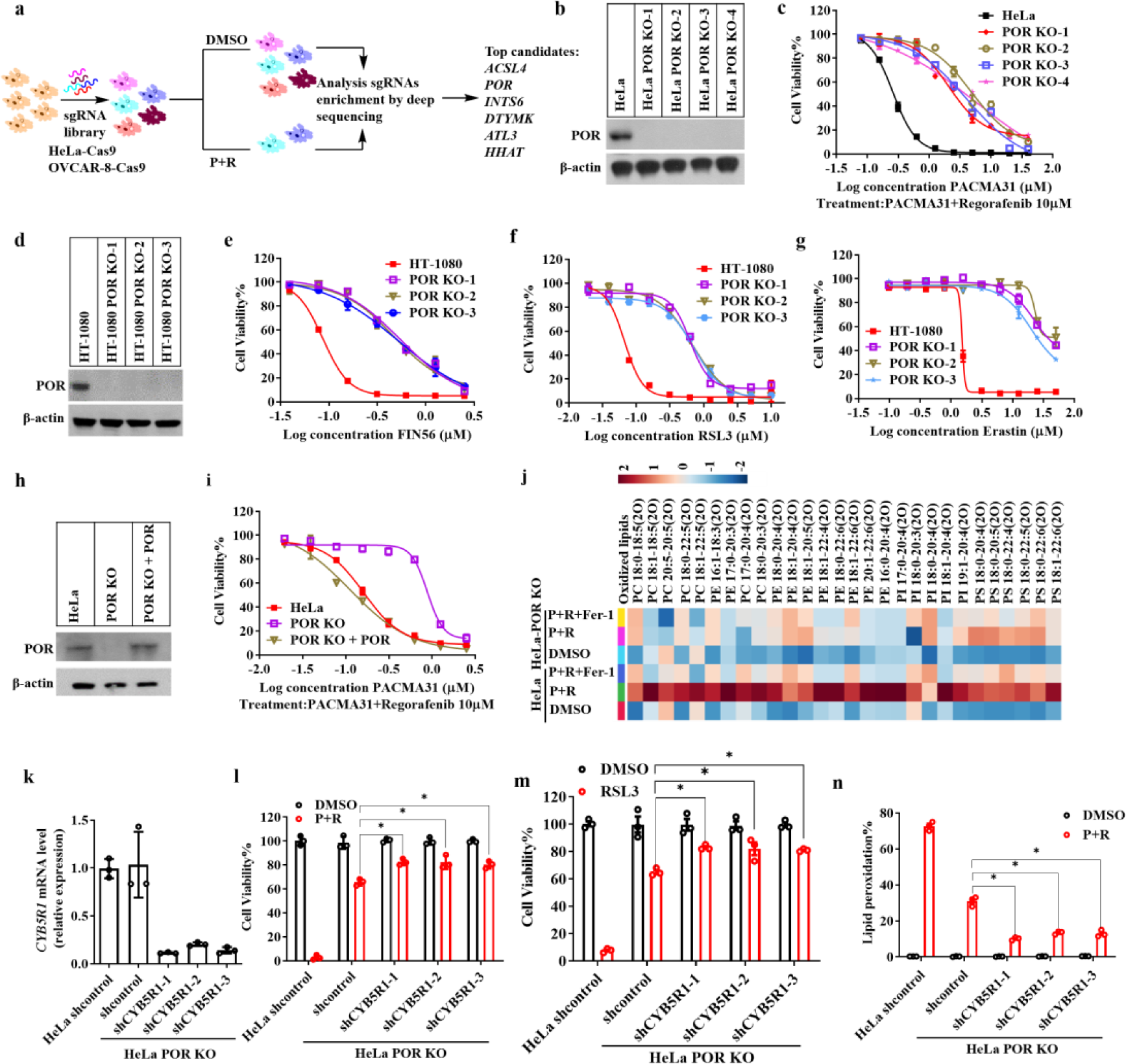
POR and CYB5R1 are required for lipid peroxidation during ferroptosis. (a) Scheme of the CRISPR-Cas9 screen strategy to identify ferroptosis regulators. sgRNAs were transduced into HeLa or OVCAR-8 Cas9 cells. P+R (PACMA31 2 μM; Regorafenib 10 μM) was used to induce ferroptosis, and the surviving cells were collected. The gRNA sequences within these cells were decoded by sequencing. (b) Immunoblot analysis of the POR protein in HeLa cells (parental or POR KO). (c) The cell viability curves in HeLa cells (parental or POR KO) treated with PACMA31 and regorafenib (P+R). (d) Immunoblot analysis of the POR protein levels in HT-1080 cells (parental or POR KO). (e-g) The cell viability curves in HT-1080 cells (parental or POR KO) treated with different ferroptosis inducers: FIN56 (e), RSL3 (f), or Erastin (g). (h) Immunoblot analysis of the POR protein levels in HeLa cells (parental, POR KO or POR KO stably expressed with POR). (i) The cell viability curves of HeLa cells (parental, POR KO or POR KO stably expressed with POR) treated with P+R. (j) LC-MS/MS-based heat maps showing peroxidized phospholipid species in HeLa cells (parental or POR KO) with the indicated treatment for 3h (PACMA31 1 μM; Regorafenib 10 μM; Fer-1 1μM). Phospholipid species in each sample (n=3) were normalized to their internal standards. (k) The relative mRNA levels of CYB5R1 in HeLa cells treated with control or shRNA targeting CYB5R1. (l-m) The cell viability of HeLa cells and HeLa POR KO cells treated with control or shCYB5R1s followed by P+R (l) or RSL3 5 μM (m). (n) The percentages of lipid peroxidation in HeLa cells and HeLa POR KO cells treated with control or shCYB5R1s followed by P+R for 1h. Lipid peroxidation was probed by Liperfluo and analyzed by flow cytometry. Cell viability throughout the figure was examined after 12h by measuring the ATP level using Cell Titer-Glo (n=3). Results were representative of three independent experiments. Student’s *t*-test (two-tailed, unpaired) was used for the comparison of the indicated two groups (mean ± SD): *p < 0.05.

POR KO cells were resistant to ferroptosis triggered by other previously characterized agents (*e.g.*, with FIN56^17^, the GPX4 inhibitor RSL3, and the xCT inhibitor Erastin^1^) as well as by GPX4 inhibition plus knockdown of FSP1 (Fig.2d-g and Extended Data Fig.4i-j). Complementation of POR expression in the POR KO cells restored sensitivity to ferroptosis-inducing agents (Fig.2h-i and Extended Data Fig.4k-l). Furthermore, we tested several commonly used apoptosis-inducing and cytotoxic chemotherapy drugs and found that none of these compounds showed decreased cytotoxicity in cells without POR (Extended Data Fig.5). Together, these findings indicated that POR is a specific participant of ferroptosis induction.

POR knockout did not impact GPX4 expression, GSH levels, or cellular phospholipid composition (Extended Data Fig.6). Instead, POR knockout prevented P+R induced lipid peroxidation measured by the fluorescence lipid peroxidation sensor Liperfluo^18^ (Extended Data Fig.7). We further performed lipidomic analysis of parental and POR KO HeLa cells and found that POR KO cells produced a significantly reduced phospholipid peroxidation compared to the parental cells after P+R administration (Fig.2j). These results indicated that POR was required for ferroptosis-related lipid peroxidation.

### CYB5R1 is required for ferroptosis

We noted that POR KO could not completely block lipid peroxidation and ferroptosis (Extended Data Fig.7), and therefore surmised that other oxidoreductases may also contribute to ferroptosis. To search for the additional participants in ferroptosis, we performed a small-scale screen on several ER-residing oxidoreductases, including *CYB5R1/2/3/4*, *FMO1/2/3/4/5*, *NDOR1*, and *ERO1B*. Upon efficient knockdown of these enzymes in HeLa POR KO cells, we found that CYB5R1, but not others, showed a significant additional protective effect in both P+R and RSL3-induced ferroptosis (Fig.2k-m and Extended Data Fig.8), with POR playing a more prominent role. Knockdown expression of CYB5R1 also significantly reduced lipid peroxidation when cells were treated with P+R (Fig.2n).

### POR’s electron acceptors are dispensable for ferroptosis

The known physiological functions of POR are quite diverse, as POR donates electrons to multiple acceptor proteins (Fig.3a)^19^. Nevertheless, its main electron acceptor partners are cytochromes P450 (CYPs), and previous studies have demonstrated that the N-terminal transmembrane domain of POR was required for its interaction with CYPs^20–22^. We thus deleted the N-terminal transmembrane domain (amino acids 2-42) of POR (Δ2-42 POR) and found that Δ2-42 POR failed to increase the CYPs activity compared with the full-length POR (Fig.3b-c). Furthermore, we incubated microsomes isolated from HeLa POR KO cells expressing either full-length POR, Δ2-42 POR, or an empty vector control alongside with four compounds (phenacetin, 17-β-estradiol, midazolam, and dextromethorphan) known to be metabolized by different CYP enzymes. Compared to cells expressing full-length POR, all compounds remained at much higher levels in HeLa POR KO cells expressing Δ2-42 POR after 8 hours of incubation (Fig.3d), indicating that deletion of amino acids 2-42 disrupted the electron transfer from POR to CYPs. However, the sensitivity to P+R induced ferroptosis was restored to a similar level by expression of either full-length POR or Δ2-42 POR in the POR KO HeLa cells (Fig.3e). This result suggested that the POR-CYPs interactions are dispensable to POR-mediated ferroptosis.

**Fig. 3.**
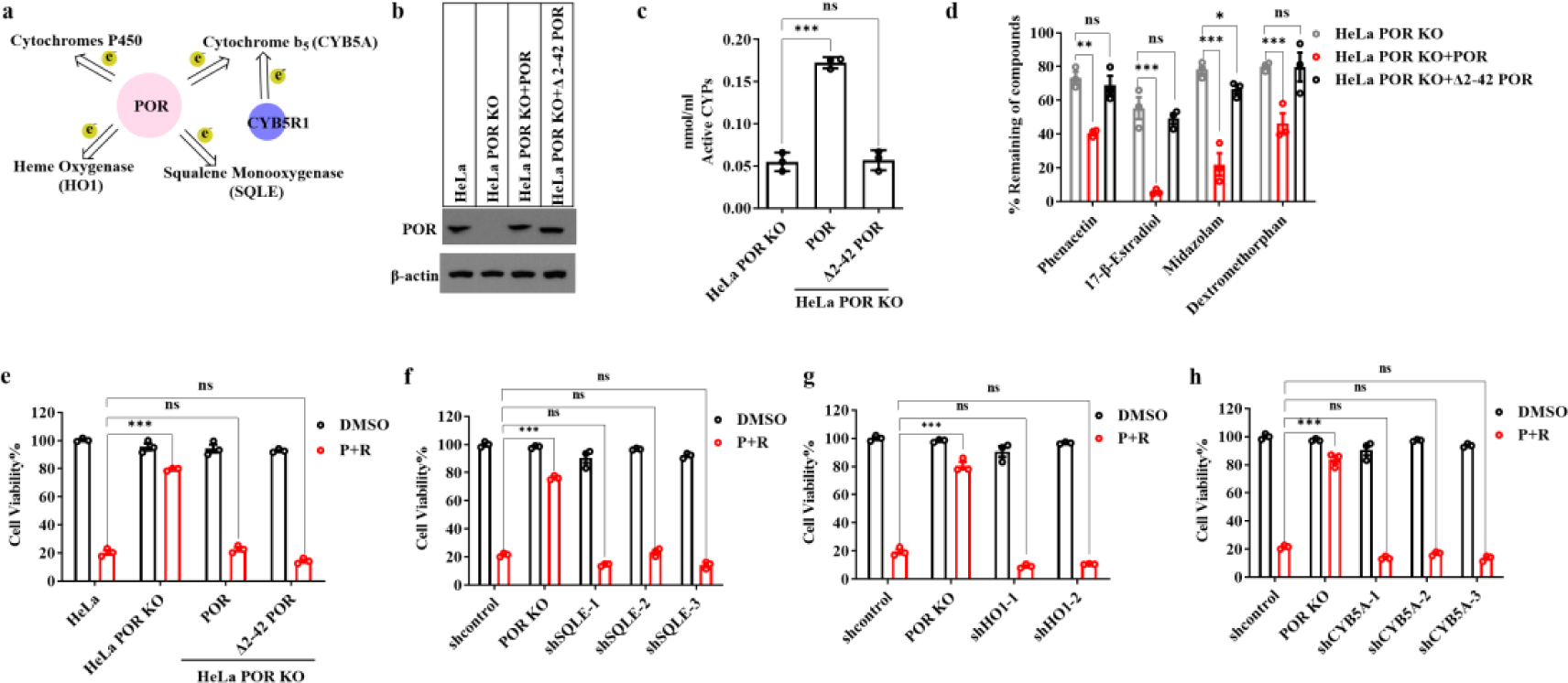
Interaction with downstream partner proteins is dispensable for POR and CYB5R1 mediated ferroptosis. (a) Scheme depicting the acceptors for electron transfer from POR and CYB5R1. (b) Immunoblot analysis of POR protein levels in HeLa and HeLa POR KO cells expressing empty vector, full length POR, or Δ2-42 POR. Δ2-42 denotes deletion of 2-42 amino acids in POR. (c) The activities of CYPs in microsomes isolated from HeLa POR KO cells expressing empty vector, full length, or Δ2-42 POR. (d) The percentages of remaining contents of indicated compounds after incubating with microsomes isolated from HeLa POR KO cells expressing empty vector, full length POR, or Δ2-42 POR for 8h analyzed by LC-MS/MS (n=3). (e) The cell viability of HeLa and HeLa POR KO cells expressing empty vector, full length POR, or Δ2-42 POR under the treatment of P+R. (f) The cell viability of HeLa (shRNA control, POR KO, or shSQLEs) cells treated with P+R (PACMA31 1 μM; regorafenib 10 μM). (g) The cell viability of HeLa (shRNA control, POR KO, or shHO1s) cells treated with P+R. (h) The cell viability of HeLa (shRNA control, POR KO, or shCYB5As) cells treated with P+R. Cell viability throughout the figure was examined after 12h by measuring the ATP level with Cell Titer-Glo (n=3, repeated in three independent experiments). Student’s *t*-test (two-tailed, unpaired) was used for the comparison of the indicated two groups with (mean ± SD): *p < 0.05; **p < 0.01; ***p< 0.001; ns, not significant.

Besides CYPs, the downstream interaction partners of POR include cytochrome b5 (CYB5A), cytochrome squalene mono-oxygenase (SQLE), and heme oxygenase-1 (HO1) ^19^. The main downstream partner of CYB5R1 is also CYB5A ^19^. Using shRNAs to knockdown the expression of CYB5A, SQLE, and HO1, we found that none of the knockdown cells exhibited resistance to P+R induced ferroptosis, indicating that these protein partners of POR/CYB5R1 also have no role in ferroptosis (Fig.3f-h and Extended Data Fig.9a-c).

### Electron transfer of POR is required for ferroptosis

Except for the aforementioned N-terminal transmembrane domain, POR included three functional domains: an FMN-binding domain, an FAD-binding domain, and an NADPH-binding domain, all critical for its electron transfer activity (Extended Data Fig.10a)^23^. We expressed a series of POR functional domain-deficient variants in HeLa POR KO cells and found that none of them conferred sensitivity to P+R induced ferroptosis (Extended Data Fig.10b-c). The requirement for the FAD-, FMN-, and NADPH-binding domains indicated that POR’s electron transfer ability mediates its function in ferroptosis.

*POR* gene mutations have been reported to cause Antley-Bixler syndrome disease, and six disease-causing allelic variants of POR all have significantly reduced electron transfer activity^24^. We expressed and purified Δ2-42 POR and the six Antley-Bixler disease mutant variants (Y178D; A284P; R454H; V489E; C566Y; V605F)-encoded enzymes from bacteria for *in vitro* enzymatic assays (Extended Data Fig.10d). None of the six variants was able to reduce cytochrome c, verifying loss of their electron transfer ability (Extended Data Fig.10e). We subsequently expressed the six mutant variants in HeLa POR KO cells and found that that none of the electron-transferring defective variants was able to restore sensitivity to P+R induced ferroptosis (Extended Data Fig.10f-g).

Upon fusing mScarlet to the C-terminus of POR (POR-mScarlet), we observed that POR is co-localized with ER-BFP-KDEL, an endoplasmic reticulum (ER) marker (Extended Data Fig.11a-c). The expression of POR-mScarlet but not a POR Y178D-mScarlet fusion protein induced generation of lipid peroxides in HeLa POR KO cells following treatment with P+R (Extended Data Fig.11c-d). Similarly, P+R treated cells expressing Δ2-42 POR-mScarlet but not Δ2-42 POR Y178D-mScarlet generated lipid peroxides extensively; the peroxides were localized at cellular membranes (Extended Data Fig.11e-f). Collectively, these results indicate that the electron transfer function of POR, but not POR localization at the ER membrane, is required for POR’s function in lipid peroxidation and ferroptosis.

### POR and CYB5R1 produce hydrogen peroxide

Since POR is a redox protein, we next explored whether its electron transfer function can result in reactive oxygen species (ROS) generation, potentially contributing to lipid peroxidation. To examine POR-mediated ROS production, we used H_2_DCFDA or hydroetidine (HE), two fluorescent probes that are respectively oxidized by hydrogen peroxide or superoxide^25–27^. Interestingly, purified Δ2-42 POR increased the levels of hydrogen peroxide (H_2_O_2_) but not superoxide (Extended Data Fig.12a-b), doing so in a manner dependent on the presence of NADPH and oxygen (Fig.4a). Inclusion of catalase—an enzyme that decomposes H_2_O_2_ to water and oxygen—inhibited oxidation of H_2_DFCDA by POR (Fig.4b), suggesting POR promotes H_2_O_2_ formation. Six variants of POR bearing mutations that have been associated with Antley-Bixler disease failed to generate H_2_O_2_ (Extended Data Fig.12c). Similar to POR, purified recombinant CYB5R1 (Δ2-28 CYB5R1) also contributed to H_2_O_2_ formation when incubated with NADH and oxygen (Fig.4c). Compared with POR, the rate of H_2_O_2_ production by CYB5R1 was much lower, probably owing to the relatively lower electron transfer efficiency of CYB5R1 (Extended Data Fig.12d-e). Moreover, we found that a combination of POR and CYB5R1 resulted in an additive effect for H_2_O_2_ production (Fig.4d).

**Fig. 4.**
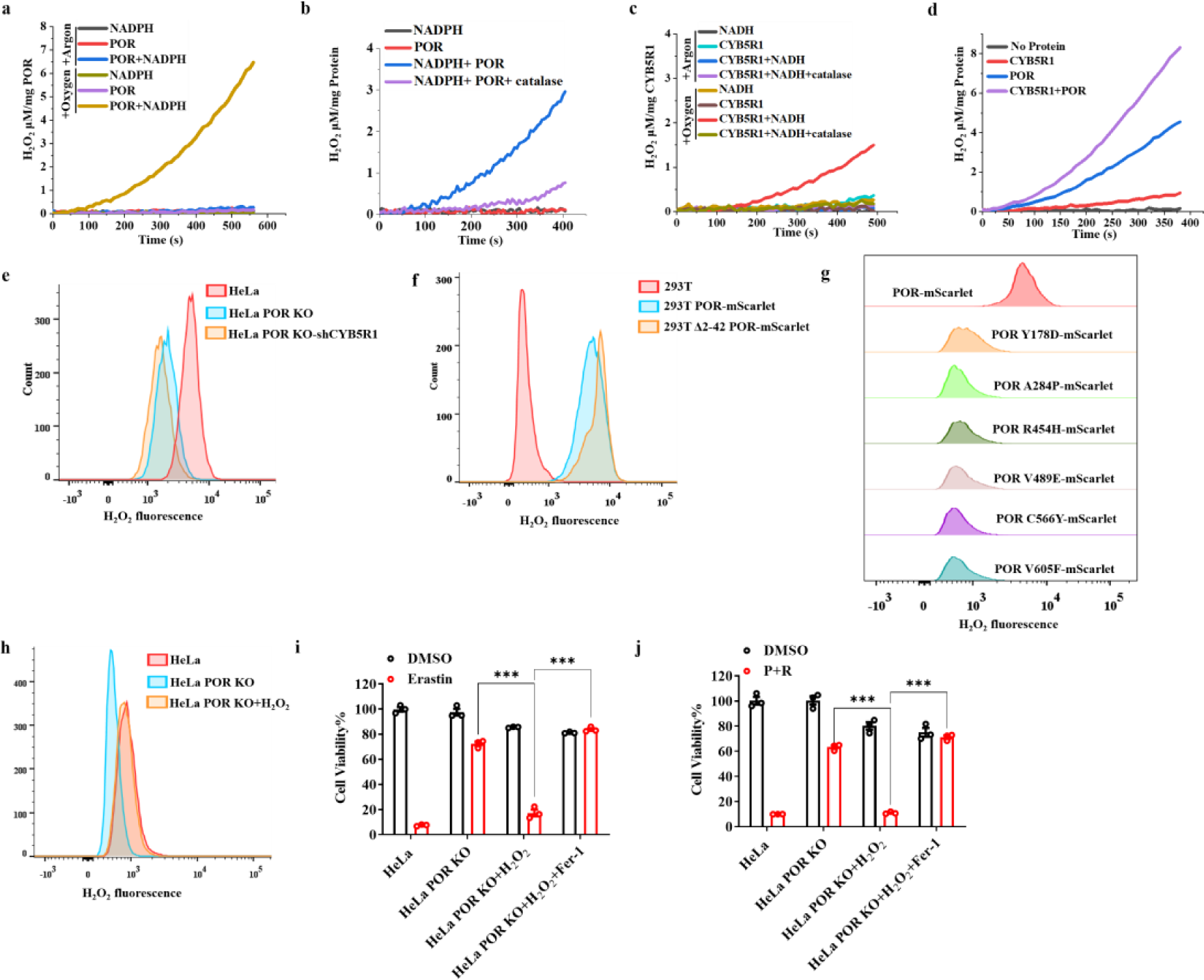
POR and CYB5R1 catalyze the generation of H_2_O_2_. (a) The H_2_O_2_ production rates of reaction buffers containing POR and NADPH in the absence (argon) or presence of oxygen. (b) The H_2_O_2_ production rates of POR in the absence or presence of catalase. (c) The H_2_O_2_ production rates of the reaction containing CYB5R1 and NADH in the absence or presence of oxygen or catalase. (d) The H_2_O_2_ production rates of the reaction containing POR, CYB5R1 alone, or both two proteins. (e) Cytosolic H_2_O_2_ production assessed in indicated HeLa cells by flow cytometry using the probe H_2_DCFDA. (f) Cytosolic H_2_O_2_ production assessed in 293T cells overexpressing POR-mScarlet or Δ2-42 POR-mScarlet by flow cytometry. (g) Cytosolic H_2_O_2_ production assessed in 293T cells overexpressing indicated POR mutants by flow cytometry. (h) Cytosolic H_2_O_2_ levels assessed in HeLa, HeLa POR KO, and HeLa POR KO cells preincubated with 30 μM H_2_O_2_ by flow cytometry. (i) The cell viability of HeLa, HeLa POR KO (incubated with H_2_O, 30 μM H_2_O_2_, or 30 μM H_2_O_2_ + Fer-1) cells treated with the xCT inhibitor Erastin. (j) The cell viability of HeLa and HeLa POR KO (incubated with H_2_O, 30μM H_2_O_2_, or 30μM H_2_O_2_+Fer-1) cells treated with P+R. Cell viability throughout the figure was examined after 12h by measuring the ATP level with Cell Titer-Glo (n=3, repeated in three independent experiments). Student’s *t*-test (two-tailed, unpaired) was used for the comparison of the indicated two groups with (mean ± SD): *p < 0.05; **p < 0.01; ***p< 0.001; ns, not significant.

Furthermore, we detected intracellular H_2_O_2_ levels by flow cytometry. Compared to HeLa cells, HeLa POR KO and HeLa POR KO-shCYB5R1 cells showed less intracellular H_2_O_2_ accumulation under normal conditions (Fig.4e). Conversely, Overexpressing POR and Δ2-42 POR in 293T cells significantly increased intracellular H_2_O_2_ levels (Fig.4f). Six variants of POR bearing mutations that reduce its electron transfer ability failed to increase intracellular H_2_O_2_ levels (Fig.4g). These results indicated that intracellular H_2_O_2_ levels are related to POR and CYB5R1 expression. However, high dose of H_2_O_2_ alone (100 μM or more) induced cell death could not be fully blocked by apoptosis or ferroptosis inhibitors (Extended Data Fig.12f). To verify that the generation of H_2_O_2_ by oxidoreductases including POR and CYB5R1 determines ferroptosis, we added different concentrations of H_2_O_2_ into culture medium and found that 30 μM H_2_O_2_ could restore the intracellular H_2_O_2_ levels of HeLa POR KO cells to that of HeLa cells (Fig.4h and Extended Data Fig.12g). The addition of 30 μM H_2_O_2_ into culture medium had no obvious cytotoxic effects and restored the sensitivity of HeLa POR KO cells to Erastin- or P+R-induced ferroptosis (Fig.4i-j). Our results suggested that H_2_O_2_ generated by oxidoreductases like POR and CYB5R1 determine ferroptosis upon GPX4/FSP1 inhibition.

### POR and CYB5R1 cause lipid peroxidation

Lipid peroxidation is usually initiated by the abstraction of hydrogen atoms from methylene carbons in PUFA. The abstraction of hydrogen atoms requires highly reactive hydroxyl radicals that were generally produced by Fenton reaction that uses ferrous iron and H_2_O_2_^28, 29^. To test if POR also mediates hydroxyl radicals generation, we used hydroxyl radical scavenger 5,5-dimethyl-1-pyrroline N-oxide (DMPO) as a hydroxyl radical probe in a POR catalyzed reaction^30^. When DMPO engages hydroxyl radicals, the process can be monitored based on specific electron spin resonance (ESR) spectra. Paramagnetic detection revealed that POR generated hydroxyl radicals when ferric chloride was included (Fig.5a). Moreover, we found that Antley-Bixler disease mutant POR Y178D failed to induce the production of hydroxyl radicals, suggesting that POR mediates hydroxyl radical generation by H_2_O_2_ production to initiate the Fenton reaction.

**Fig. 5.**
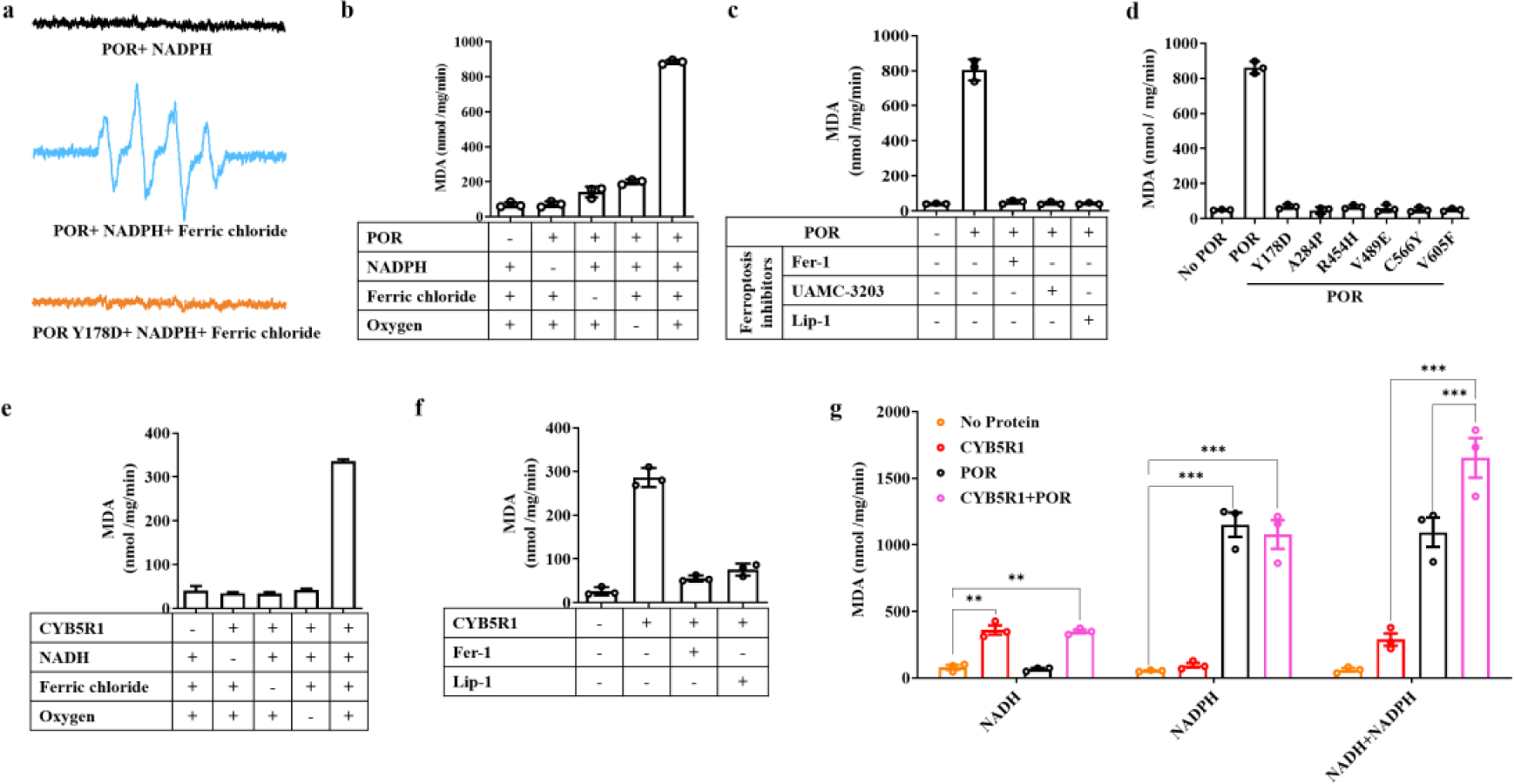
POR and CYB5R1 catalyze the generation of lipid peroxidation. (a) The ESR spectrums of hydroxyl radicals generated by POR in the presence of oxygen. (b) Lipid peroxidation catalyzed by POR was evaluated by measuring the MDA content of the indicated reaction mixtures. (c) The production of MDA in the presence of POR with different ferroptosis inhibitors (50 μM each). (d) The production of MDA in the presence of different POR variants (100 nM each). (e) The production of MDA in the presence of CYB5R1 (100 nM). Ferric chloride (120 μM), NADH (200 μM), oxygen, and phospholipid mixture (100 μg/ml) were also included in the reaction buffers. (f) The production of MDA in the presence of CYB5R1 with different ferroptosis inhibitors. (g) The production of MDA in the presence of POR, CYB5R1 alone, or both proteins. The data were representative results of three independent experiments. Bar graphs show mean ± SD.

Based on the above observation, we next tested if POR could cause lipid peroxidation in a reaction with defined components. To this end, we used a phospholipid mixture as a substrate for thiobarbituric acid (TBA) assays to measure lipid peroxidation. Malondialdehyde (MDA) is a major degradation product of lipid hydroperoxides, and MDA can react with two molecules of TBA to form a stable complex that can be quantified using spectrophotometry^31^. We found that POR catalyzed lipid peroxidation when phospholipids were used as substrates; Ferric chloride, oxygen, and NADPH were also essential (Fig.5b). Moreover, we found that the extent of POR-catalyzed lipid peroxide formation increased in a dose-dependent manner upon increases in ferric chloride and NADPH concentrations (Extended Data Fig.13a-b). We also tested other transition metal ions and found that iron was the only one that functioned in POR catalyzed lipid peroxidation (Extended Data Fig.13c). When we added the ferroptosis inhibitors (*e.g.*, the lipid radical scavenger Fer-1, UAMC-3203^32^, and Lip-1) in the enzymatic assays, POR-catalyzed lipid peroxidation was significantly inhibited (Fig.5c). Furthermore, the six Antley-Bixler disease mutant POR variants failed to catalyze lipid peroxidation (Fig.5d).

Similar to POR, CYB5R1 catalyzed the formation of lipid peroxides in the presence of oxygen, ferric ions, and NADH (Fig.5e). We also tested other transition metal ions and found that iron was also specific in CYB5R1-catalyzed lipid peroxidation (Extended Data Fig.13d). When the ferroptosis inhibitors (e.g., the lipid radical scavengers Fer-1 and lip-1) were included in the reaction, CYB5R1-catalyzed lipid peroxidation was significantly reduced (Fig.5f). Furthermore, we found that the CYB5R1-catalyzed lipid peroxidation was less efficient than POR (Fig.5g), which was consistent with its lower H_2_O_2_ production compared with POR (Fig.4d). When combined, POR and CYB5R1 induced more robust lipid peroxidation (Fig.5g). These experiments indicated that CYB5R1 works together with POR in ferroptosis-related lipid peroxide formation.

### POR and CYB5R1 induce liposome rupture

Lipid peroxidation and its attendant membrane damage is the signature of ferroptosis^7^; however, the detailed mechanism(s) for how lipid peroxidation leads to membrane rupture remain unknown. We used purified recombinant POR/CYB5R1 and the other required components for lipid peroxidation (NADPH/NADH and iron) to test whether the lipid peroxidation mediated by these specific components is sufficient to cause leakage from liposomes. We prepared liposomes with a phospholipid mixture and enclosed Tb_3+_ ions, followed by incubation with purified POR, NADPH, and ferric chloride. Tb_3+_ normally has weak fluorescence; however, after release from liposomes and binding to DPA, the fluorescence signal of Tb_3+_/DPA chelate increases ∼100-fold, yielding a strong signal representing liposome leakage^33^. We found that more than 60% of the Tb_3+_ in liposomes was released within 40 min in assays that included POR, ferric chloride, and NADPH during (Fig.6a). Further, this liposome leakage was significantly blocked by the presence of the ferroptosis inhibitors Fer-1 and Lip-1 (Fig.6b), indicating that lipid peroxidation can account for the observed liposome leakage. None of the six aforementioned electron-transfer defective Antley-Bixler disease mutant POR variants caused liposome leakage (Extended Data Fig.14a).

**Fig. 6.**
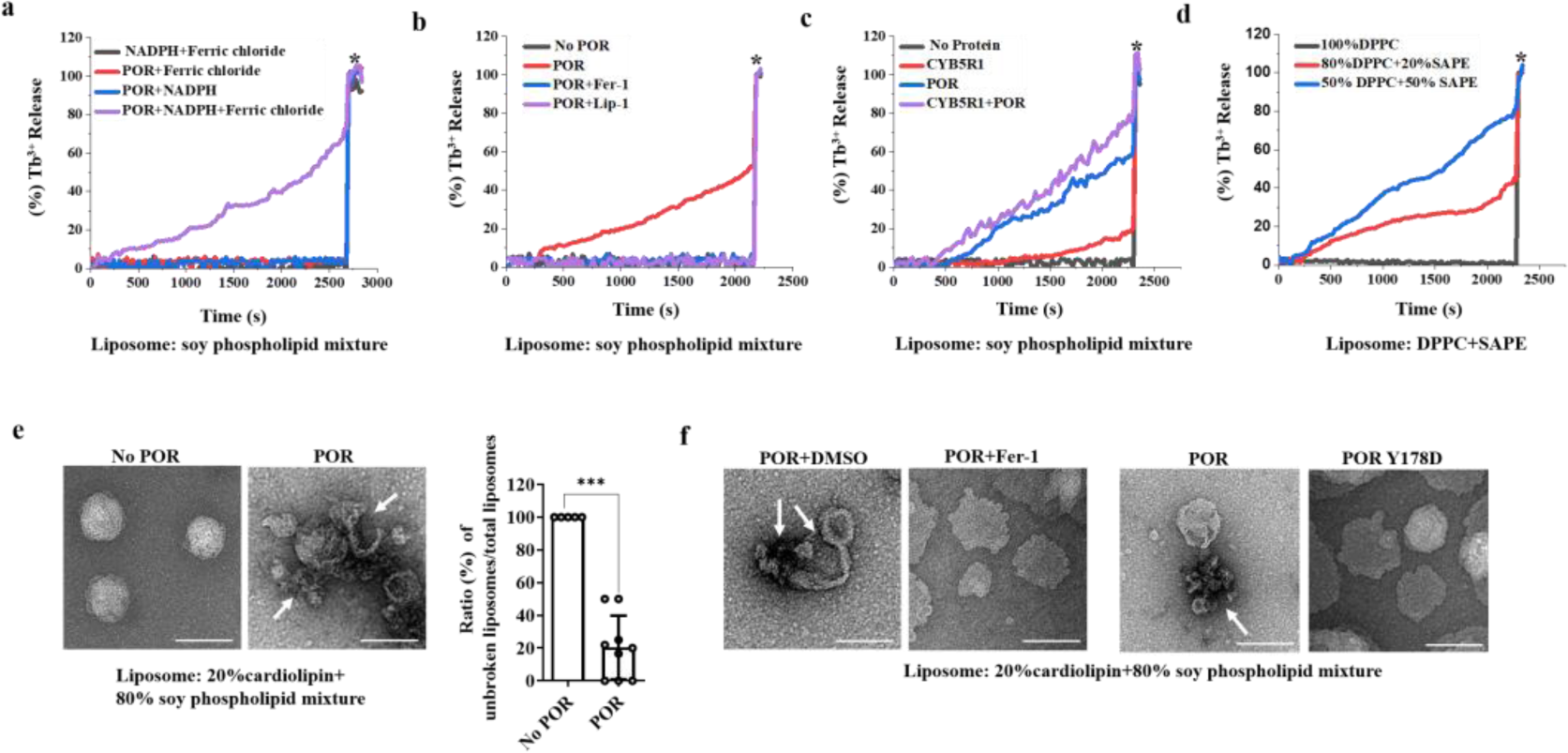
POR and CYB5R1 cause liposome leakage. (a) Time courses of liposome leakage. Asterisks indicated the time points when Triton X-100 was added to achieve complete release of Tb_3+_. (b) Fer-1 and Lip-1 blocked POR induced liposome leakage. (c) The liposome leakage caused by POR, CYB5R1, or both proteins. (d) The leakage of different ratios of SFA-phospholipids SAPE and PUFA-phospholipids DPPC containing liposomes caused by POR. (e) Negative-stain electron microscopy of liposomes treated with or without POR. White arrowheads indicated broken liposomes. The ratios of unbroken liposomes/ total liposomes were calculated (n>40 of liposomes in each group). Scale bars, 200 nm. Student’s *t*-test (two-tailed, unpaired) with mean ± SD was used for the comparison of the indicated two groups: ***p< 0.001. (f) Transmission electron microscopy of liposomes treated with POR or POR Y178D in the absence or presence of Fer-1. All data were representative of three independent experiments.

Similar to POR, CYB5R1 also caused liposome leakage, albeit with much reduced efficiency. In the CYB5R1 assays, 20% of Tb_3+_ was released from liposomes within 40 min (Fig.6c). The combination of POR and CYB5R1 showed an additive effect on liposome leakage: 80% of the Tb_3+_ was released within 40 min (Fig.6c). Besides these assays with liposomes comprising a phospholipid mixture, we assayed cardiolipin-containing liposomes and found that the same amount of POR caused 100% Tb_3+_ leakage from these cardiolipin-containing liposomes (Extended Data Fig.14b). We speculate that this increased level of liposome leakage may related to the high PUFA content of cardiolipin. Supporting this, we prepared liposomes containing different ratios of the PUFA-phospholipid SAPE (1-octadecanoyl-2-(5Z,8Z,11Z,14Z-eicosatetraenoyl)-sn-glycero-3-phosphoethanolamine) and the saturated fatty acid (SFA)-phospholipid DPPC (1,2-dipalmitoyl-sn-glycero-3-phosphocholine), and assays showed that POR was able to release about 80% of Tb_3+_ from 50% SAPE containing liposomes while only 40% release was observed with the 20% SAPE containing liposomes. Finally, we found POR was not able to break liposomes comprising 100% SFA (Fig.6d), revealing that POR-mediated liposome leakage is PUFA-phospholipid dependent.

We also performed transmission electron microscopy to directly observe morphology changes in liposomes treated with POR. Compared to liposomes without POR treatment, incubation of POR (with NADPH and ferric chloride) disrupted the phospholipid membranes and caused obvious rupture of liposomes (Fig.6e). Addition of Fer-1 inhibited POR-induced liposome rupture, and we found that the Y178D POR variant (dysfunctional electron transfer capacity) failed rupture liposomes (Fig.6f). Note that these electron microscopy results were consistent with corresponding Tb_3+_ leakage experiments (Extended Data Fig.14c-e). Collectively, these results demonstrate that lipid peroxidation catalyzed by POR disrupts PUFA-containing phospholipid membrane integrity.

### POR knockdown conferred protection during acute liver injury

Our cellular level studies prompted further analysis of POR functions in ferroptosis-related pathology *in vivo*. Consistent with previous studies indicating that ferroptosis was involved in acute liver injury (ALI)^34, 35^, survival analysis revealed that the ferroptosis inhibitor UAMC-3203^32^ protected mice from lethal doses of concanavalin A (Con A) (Fig.7a). Con A is a T-cell mitogenic lectin derived from the jack-bean plant and is general used as an *in vivo* model to mimic autoimmune hepatitis (AIH), an immune-mediated ALI caused by chronic autoimmune-driven attack of liver cells that leads to the destruction of hepatic parenchyma^36^. To examine whether POR knockdown protects cells from ALI by inhibiting ferroptosis upon Con A exposure, we first designed a specific shRNA targeting POR mRNA for *in vivo* administration via the adeno-associated virus (AAV) and verified that AAV infection was capable of significantly downregulating POR expression in the liver. Then, we treated mice with lethal doses of Con A to induce ALI. As expected, 70% of shPOR mice survived whereas 100% of vehicle-treated shRNA control mice died within 24 h after application of Con A (Fig. 7b). Notably, shPOR mice exhibited a relatively lower protective effect than UAMC-3202 (Fig.7a-b), which was consistent with our cell assay results indicating that other oxidoreductases were also involved in ferroptosis.

**Fig. 7.**
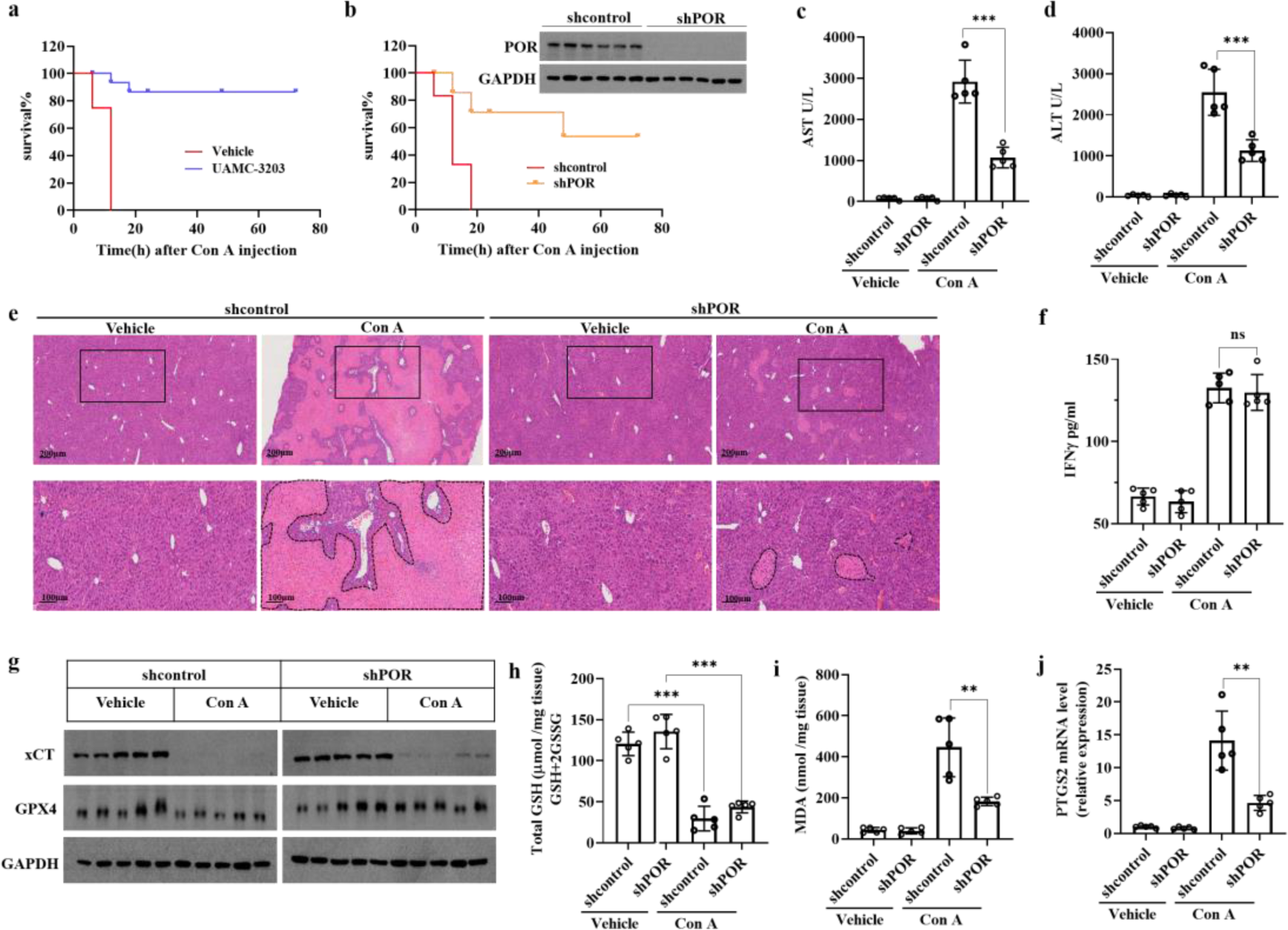
The knockdown of POR *in vivo* attenuates concanavalin A-induced acute liver injury. (a) Analysis of mice survival upon treatment with a lethal dose of concanavalin A (Con A, 30 mg/kg) using the Kaplan–Meier method (n = 8 mice each group). Mice treated with vehicle or ferroptosis inhibitor UAMC-3203 (10 mg/kg) 1 h prior to injection of Con A. (b) Analysis of shRNA control or shPOR mice survival upon treatment with a lethal dose of concanavalin A (Con A, 30 mg/kg) using the Kaplan–Meier method (n=6 mice each group). The POR expression level in the livers of shRNA control or shPOR mice were displayed at the top right corner. (c-d) Serum levels of AST (c) and ALT (d) were assessed in shRNA control or shPOR mice treated with vehicle or 15 mg/kg Con A (n = 5 mice each group). Serum samples were obtained 24 h after Con A injection. (e) Hematoxylin and eosin (H&E) staining in liver cross sections from vehicle or Con A-treated mice (dashed lines show necrosis). Insets show respective picture. (f) Serum levels of IFN-γ in mice measured by ELISA. (g) The expression of xCT and GPX4 in liver tissues of indicated groups. (h) Total GSH levels in livers of indicated groups. (i) The MDA levels in livers of indicated groups. (j) The relative mRNA expression of PTGS2 in livers of indicated groups. The relative PTGS2 levels in the livers of shRNA control mice treated with vehicle was set to 1.0. Bar graphs show the mean ± SD. Student’s t test (two-tailed, unpaired) was used for the comparison of the indicated two groups: *p < 0.05; **p < 0.01; ***p< 0.001; ns, not significant.

To further study the potential role of POR in ALI, shRNA control or shPOR mice were injected with sublethal doses of Con A to induce liver injury (Extended Fig.15a). Serum levels of ALT and AST were significant elevated at 24 h after Con A injection in shRNA control mice, and the degree of up-regulation of ALT and AST were much lower in shPOR mice (Fig.7c-d). Consistently, histological analyses indicated that POR knockdown significantly decreased necrotic tissue area in liver than compared to the Con A-treated shRNA control group (Fig.7e). During the pathological process of AIH, several proinflammatory cytokines are increased^37^. Indeed, Con A induced significant production of IL4, TNFα, and IFNγ in early pathology stages and there was no difference among shRNA control and shPOR mice (Fig.7f and Extended Data Fig.15b-c). A previous study showed that IFNγ can downregulate the expression of xCT to induce ferroptosis in tumor cells^38^. Thus, we determined the expression of the key ferroptosis regulators xCT and GPX4 in liver after Con A treatment. As expected, both shPOR and shRNA control group showed significantly lower expression of xCT when treated with Con A. GPX4 expression levels showed no obvious change (Fig.7g). Consistent with the decreased expression of xCT, the hepatic levels of GSH were clearly decreased upon Con A injection, and the level of GSH depletion was similar in shRNA control and shPOR mice (Fig.7h and Extended Data Fig.15d-e). To detect if POR-mediated lipid peroxidation acted downstream of GSH depletion in ALI, we assessed the content of the lipid peroxidation marker MDA and mRNA expression levels of PTGS2^3^, which is also a key marker of lipid peroxidation and ferroptosis *in vivo*. Compared to shRNA control group, shPOR mice showed much lower MDA and PTGS2 levels when treated with Con A (Fig.7i-j). Together, our *in vivo* studies strongly suggest that Con A-induced reduction of xCT and the attendant GSH levels caused ferroptosis in mouse liver. Moreover, knockdown of POR could not block either Con A-induced elevation of cytokines, or IFNγ-induced reduction of xCT/GSH levels, blocked liver cell ferroptosis, and therefore blocked Con A-induced liver damage and animal death.

## DISCUSSION

Although the NADH–FSP1–CoQ10 and glutathione–GPX4 systems are well-characterized in their ability to suppress ferroptosis, the nature of any mechanism(s) underlying the iron-dependence in ferroptosis-related accumulation of lipid peroxides remains unclear. In the present study, facilitated by a robust ferroptosis stimulation assay comprising P+R co-treatment, we discovered that ER-residing oxidoreductases including POR and CYB5R1 are the true “executioners” for ferroptosis. Unlike a recent report that also identified POR as a required component for ferroptosis^39^, we demonstrated that POR interaction with downstream partners (including CYPs/HO-1/SQLE/CYB5A) is irrelevant to ferroptosis, since disruption of interactions between POR and CYPs or knockdown POR or CYB5R1 downstream proteins did not affect ferroptosis. Moreover, we proved that the function of these oxidoreductases in ferroptosis is an “incidental” donation of electrons from NADPH/NADH to oxygen to generate hydrogen peroxide, which reacts with iron in a Fenton reaction to produce highly reactive hydroxyl radicals that react with PUFA on the membrane to carry out lipid peroxidation (Extended Data Fig.16). Therefore, our results indicated that it is oxidoreductases, which generate hydrogen peroxide under normal conditions, that account for ferroptosis when the protective antioxidant system is inhibited. As double inhibition of POR and CYB5R1 still could not fully prevent ferroptosis in cells, it is tempting to speculate that other oxidoreductases, like the NOXs1, in some specific situations ALOXs40, or others, might also additively contribute to ferroptosis due to their role in generating hydrogen peroxide under normal conditions in cells.

Unlike other regulated cell death programs elucidated so far, including apoptosis, necroptosis, and pyroptosis, wherein the cell death programs must be “turned on” by specific physiological stimuli, ferroptosis seems to be constitutively “on”, and must be actively suppressed by the GPX4/FSP1 lipid peroxidation reduction systems. Since redox reactions are involved in many biological processes, the reduction or defection of GPX4/FSP1 systems, either in diseases or aging, would result in ferroptotic cell death. Conversely, specific pharmacological inhibition of these two lipid reduction systems in cancer cells may also be an attractive therapeutic strategy.

Notably, pathophysiological roles for ferroptosis in human diseases such as tumor suppression, neurodegeneration, and organ damage are extensively documented. More and more studies have revealed that ferroptosis is a significant contributor to tissue injury in the liver and kidney^2^. We found that POR knockdown in the liver indeed inhibits ferroptosis and protected against immune-mediated acute liver injury, thereby indicating that specifically targeting POR could be further evaluated as an innovative therapeutic strategy for treating ferroptosis-implicated diseases.

## Methods

### Cell culture

All mammalian cells were cultured at 37°C with 5% CO_2_. HCT-116, HT-1080, HeLa, MEF and 293T cells were cultured in DMEM (GIBCO) and OVCAR-8 was cultured in RPMI-1640 (GIBCO), both DMEM and RPMI were supplemented with 10% FBS (Invitrogen) and 1% penicillin and streptomycin.

### Cell viability assay

Cell viability assays were performed by measuring the cellular ATP level using a Cell Titer-Glo Luminescent Cell Viability Assay kit (Promega) according to the manufacturer’s instructions. The luminescence intensity was read by a microplate reader (PerkinElmer EnSpire). All data were analyzed using GraphPad Prism (GraphPad Software, Inc., San Diego, CA, USA). The curves were fitted using a non-linear regression model with a sigmoidal dose response.

### Western blotting

Cells were lysed either in RIPA-lysis buffer (50 mM Tris-HCl pH 7.4, 150 mM NaCl, 1% TritonX-100, 1% sodium deoxycholate, 0.1% SDS and cOmplete™ EDTA-free Protease Inhibitor Cocktail (Roche)) or directly solubilized in 1x sodium dodecyl sulfate (SDS) buffer, then subjected to SDS-polyacrylamide gel electrophoresis (SDS-PAGE). Cells lysed with RIPA-lysis buffer were set on ice for 30 min, and then centrifuged at 20,000 ×g for 10 minutes. The supernatants were then mixed with 4x SDS buffer and subjected to SDS-PAGE. The following antibodies were used in western blotting: Anti-POR Rabbit mAb (Abcam, Cat# ab180597); Anti-GPX4 Rabbit mAb (Abcam, Cat# ab125066); Anti-β-actin Rabbit mAb (Cell Signaling Technology, Cat# 4970); Anti-Rabbit HRP (Sigma-Aldrich, Cat# A0545); Anti-Mice HRP (Sigma-Aldrich, Cat# A9044); Anti-Flag HRP (Sigma-Aldrich, Cat# A8592); and Anti-His-HRP (Miltenyi Biotec, Cat# 130-092-785).

### Virus packaging and transfection of shRNAs

The shRNA-containing pLKO.1 lentiviral plasmids (Sigma-Aldrich) were co-expressed with pMD.2G and psPAX2 in 293T cells for 48 hours with Lipofectamine 3000 (Thermo Fisher) according to the manufacturer’s instructions. Supernatant (containing virus particles) was filtered through a 0.22 µM filter and used to infect cells which were seeded into six-well plates at a density of 1 × 10^5^ cells per well with polybrene (5 µg/mL). Cells were selected with puromycin 48 h after viral transduction before use in experiments. The shRNAs used were from Sigma shRNA clone library which were identified by TRC numbers.

### RT-qPCR

Total RNA was prepared using the TRIzol^TM^ Reagent (Invitrogen) according to the manufacturer’s instructions. After reverse transcription, RT-qPCR was performed using Takara SYBR Premix Ex Taq kits to detect the mRNA expression levels of indicated genes. The expression levels of target genes were normalized by subtracting the corresponding β-actin/GAPDH threshold cycle (CT) value.

### Construction of HeLa/OVCAR-8 Cas9 cells

HeLa/OVCAR-8 cells were seeded in a six-well plate (1×10^5^ cells/per well) for 12 h. Each well was infected with the Cas9 lentivirus (2 ml) for 24 h and then fresh medium was replaced in the wells. After three days, the GFP-positive cells were sorted using a BD FACSAria Fusion flow cytometer to select single clones of HeLa/OVCAR-8-Cas9 cell lines. The single clones were amplified and immunoblotting was performed to select positive clones for further experiments.

### Genome-wide CRISPR-Cas9 screen

The lentiviral gRNA library for the CRISPR-Cas9 screen was obtained commercially from Addgene (#50947); The library was amplified, and the lentivirus was prepared following the manual instructions. To perform the genome-wide screen, HeLa-Cas9 cells were seeded in a 15 cm dish (1×10^7^ cells) and infected with the gRNA lentivirus library with an MOI of 0.3. At 24 h after infection, 10 mg/ml puromycin was added into dishes to kill uninfected cells. Then the infected cells were amplified and seeded into 15 cm dishes (1×10^7^ cells per dish). Three dishes were combined as one group and treated with PACMA31 (2 μM) and regorafenib (10 μM) to trigger ferroptosis; another group were treated with DMSO as control. The screens were performed in HeLa and OVCAR-8 cas9 cells and repeated twice. The surviving cells after ferroptosis induction were collected and lysed to extract whole genomic DNAs. The DNAs were further precipitated and re-dissolved in ddH_2_O as the templates for amplification of the gRNA.

The gRNAs were amplified by the PCR strategy. Final PCR products were subjected to electrophoresis and sequenced using a HiSeq2500 instrument (Illumina). The first 19 nucleotides from the sequencing read were the gRNA sequence recovered from the library. The frequency of each gRNA was obtained by dividing the gRNA read number by the total sample read number; the fold enrichment was calculated by comparing the frequency of each gRNA in the P+R treated group with that in the control group.

### The construction of CRISPR-Cas9 POR knockout cells

pX458-POR plasmid was transfected into cells seeded in six-well plates using the transfection reagent (Lipofectamine 3000, Thermo Fisher) following the manufacturer’s instructions. After two days, the GFP-positive cells were sorted using a BD FACSAria Fusion flow cytometer to select single clones. The single clones grew in 96 well-plates for a week and were then transferred to 24 well-plates; further immunoblotting was conducted to select positive clones. The gRNA was designed as follows: POR-gRNA-1 (GCTCCACCGTGTCCGAGGCGG); POR-gRNA-2 (GATTCTGTTTTCGCTCATC G); POR-gRNA-3 (GAGGAACATCATCGTGTTCTA); POR-gRNA-4 (GAGGCAT GTCAGCGGACCCTG).

### Protein re-expression in HeLa POR KO cells

Using the full-length cDNAs for human POR as a template, we amplified the DNA fragments of POR (full-length and truncates), then the fragments were inserted into pWPI vector to generate the pWPI-POR (full-length and truncates) constructs. Mutants were generated by a quick mutagenesis kit (Genestar) and verified by Sanger sequencing. The plasmids of pWPI-POR (WT, mutants and truncates) were co-expressed with pMD.2G and psPAX2 into 293T cells for 48 hours with Lipofectamine 3000 (Thermo Fisher) according to the manufacturer’s instructions. Virus containing supernatant were filtered and used to infect cells that were seeded into six-well plates (1 × 10^5^ cells per well) with 5 µg/mL polybrene. After three days, the GFP-positive cells were sorted using a BD FACSAria Fusion flow cytometer to select single clones. The single clones were grown in 96-well plates for at least 1 week then transferred to 24-well plates to detect the protein expression levels and select positive clones for further experiments.

### Compound screen assay

Compound screens were performed at the Chemistry Center at National Institution of Biological Science, Beijing. OVCAR-8 cells (2,000 cells per well) were seeded in a 384-well plate on day one. On day two, about 300 compounds from the FDA-approved drug sub-library were individually added in each well to a final concentration of 10 µM. Then, 500 nM PACMA31 was added into each well. Cell viability was assessed 12 hours after treatment. Hits with significant reduction in cell viability were selected for further verification.

### Pull down assay of PACMA31-Probe

OVCAR-8 cells were seeded in triplicate in 10 cm dishes one day before the start of the experiment. Cells were incubated with DMSO, or PACMA31 (1 μM), or RSL3 (1 μM) for 1 h; the PACMA31-Probe (1 μM) was then added and the samples were incubated for an additional 1 h. The cells were collected and lysed in RIPA lysis buffer. Subsequently, cell lysates were incubated for 2 h for the click chemistry reaction after addition of Biotin-N_3_ (10 μM), CuSO_4_ (50 μM), Tris [(1-benzyl-1H-1,2,3-triazol-4-yl) methyl] amine (TBTA, 50 μM), and tris(2-carboxyethyl) phosphine (TCEP, 50 μM). The proteins were precipitated by adding 5 sample volumes of methanol and centrifuging at 20,000 ×g for 5 min. Precipitated protein was washed once with methanol and the pellet was re-suspended in PBS containing 2% SDS and 10 mM DTT. Once dissolved, the samples were diluted 10-fold with PBS to 1 mg/mL and 0.2% SDS. The biotin-modified proteins were then enriched using streptavidin beads; the beads were washed three times with PBS containing 0.2% SDS. The pull-down fraction was eluted in 1x protein loading buffer for 15 min at 95°C, followed by immunoblotting with antibodies against GPX4 and β-actin.

### Assessment of lipid peroxidation using Liperfluo

A total of 1 x 10^5^ cells per well were seeded on six-well plates one day prior to the experiment. On the next day, cells were treated with the ferroptosis inducers, then cells were incubated with Liperfluo (DojinDo) 1 μM for 30 min at 37°C before they were harvested by trypsinization. Subsequently, cells were resuspended in fresh PBS strained through a 70 μM-cut off cell strainer and analyzed using the 488 nm laser of a flow cytometer for excitation. Data was analyzed using FlowJo Software.

### Determination of intracellular GSH levels

Cells were seeded on 10 cm dishes (1×10^6^ cells per dish) and cultured for 24 h. Tissues were lysed by 5% Total glutathione content was measured using an Intracellular glutathione (GSH) Detection Assay Kit (Abcam) as described in the manufacturer’s protocol. Protein quantification was performed using the Pierce BCA Protein Assay Kit (Thermo Scientific) as described in the manufacturer’s guidelines.

### Protein expression and purification

The PCR products of POR (Δ2-42 WT or mutants) or CYB5R1 (Δ2-28) were subcloned into pET28a. Proteins were expressed in *E. coli* strain JM109 (DE3) cultured in Luria-Bertani (LB) medium containing 100 μM riboflavin and 50 μg/ml kanamycin. The bacteria were cultured at 37°C to reach the OD value of 0.8, then 1 mM isopropyl β-D-1-thiogalactopyranoside (IPTG) was added into LB to further culture overnight at 15°C. The bacteria were harvested by centrifugation for 30 min at 4,000 ×g and lysed in the presence of cOmplete™ Protease Inhibitor Cocktail (Roche) and Phenylmethylsulfonyl fluoride (PMSF). The supernatant was isolated by centrifugation for 30 min at 20,000 ×g and then incubated with a Ni_2+_-NTA column (QIAGEN) for 5h. Then the columns were washed and eluted with 300 mM imidazole. The proteins were concentrated and analyzed by SDS-PAGE.

### Confocal microscopy

Cells were seeded in 35 nM glass-bottom culture dishes (Mat Tek), then treated with the compounds indicated in the relevant figure captions. Lipid peroxidation was detected by Liperfluo (DojinDo); Liperfluo was added into the medium for 15 min. Subsequently, cells were washed twice with PBS and live cell images were acquired using a Nikon A1 microscope.

### Microsomes isolation

The cells were seeded in 15 cm dishes and were collected from ten dishes at a time as a group. The cell pellets were mixed with 10-fold volume of sucrose buffer (0.25 M sucrose; protease and phosphatase inhibitor cocktails (Roche)) and homogenized until a completely homogenous solution was obtained. The solution was centrifuged at 10,000 ×g for 10 min at 4°C. The supernatants were further isolated by centrifugation at 30,000 ×g for 90 min. The remaining pellet was mainly microsomes. The microsomes were resuspended and diluted according to the protein concentration measured in the supernatants using the Pierce BCA Protein Assay Kit (Thermo Scientific).

### Measurement of CYPs in microsomes

The microsomes were diluted to the same protein concentrations in different groups with 100 mM of potassium phosphate buffer (pH 7.5) containing 1.0 mM of EDTA, 20% glycerol (vol/vol), 0.5% sodium cholate (wt/vol) and 0.4% (wt/vol) Triton N-101. A total of 1 mM NADPH was added into the buffer and incubated for 1 h, then each sample was divided into two groups, one group was set as the control, while the other group was slowly bubbled with CO for 5 min to completely induce the interaction of CO with iron (II) in P450s. Both groups were placed in a spectrophotometer (Tecan GENios) to record the absorbance between 400 and 500 nm. The activity of P450s were calculated as follows: [(A_450_ – A_490_) – (A_450_ – A_490_) _control_]/0.091 = nmol P450 of per ml.

### Measurement of the drug metabolism ability of CYPs in microsomes

Microsomes resuspended in PBS buffer were incubated with 100 μM of a mixture containing phenacetin, 17-β-estradiol, midazolam, and dextromethorphan. The mixture was pre-incubated for 5 min at 37°C, then NADPH was added into the reaction solution to the final concentration of 1 mM. The mixture was then vortexed gently and 20 µL was immediately aliquoted after incubation and set as 0 h. After 8 h of incubation at 37°C, 20 µL was removed from the reaction buffer. The 0 h and 8 h aliquots were added into 300 µL methanol to quench the reaction. The samples were vortexed vigorously for 1 min, then placed at −20°C for 1 h. Samples were centrifuged at 3,500 g for 15 min and 100 µL of the supernatant was analyzed using LC-MS/MS. The 0 h mass response was set to 100%.

### Measurement of the electron transfer ability of POR in NADPH–cytochrome c reduction activity

Horse heart cytochrome c (Sigma) and Δ2-42 POR were dissolved in potassium phosphate buffer (10 mM, pH 7.7) to final concentrations of 50 μM and 100 nM, respectively. The basal absorbance at 550 nm was recorded using a microplate reader (Tecan GENios). Then the 200 mM NADPH stock solution was added to the final concentration of 200 μM to initiate the reaction and the absorbance was immediately recorded at 550 nm and then every 5s until the value did not change anymore. The activity of POR was calculated according to the reduction of cytochrome c: (ΔA_550_/min)/0.021= nmol of cytochrome c reduced per min.

The electron transfer ability of Δ2-28 CYB5R1 was performed similar with Δ2-42 POR, except that NADPH was replaced by NADH.

### Lipidomic analysis

HeLa cells (WT or POR KO) were seeded into 15 cm dishes and three dishes were recognized as one sample; three replicates were used. HeLa cells (WT or POR KO) were treated with DMSO, P+R or P+R+Fer-1 for 2 h. Then cells were washed twice using PBS, trypsin digested, and collected in centrifuge tubes. The cells were further washed twice with PBS and centrifuged at 1000 ×g for 3 min; the supernatant was discarded. The cell pellets were stored at −80°C until the samples were examined. For peroxided lipidomic analysis, the samples were examined immediately after the cells were collected. Lipids were extracted by adding a 1 ml mixture of chloroform and methanol (2:1) and 10 µl lipid mixture of internal standard (Avanti) into each sample; samples were then sonicated for complete extraction. A total of 100 μl H_2_O was added into the solution to promote stratification of the extraction solutions. The samples were vortexed vigorously and centrifuged at 15,000 ×g for 5 min. The chloroform layer was removed to a new tube and vacuum dried. A total of 100 μl methanol was added to the samples to resolve the lipids, centrifuged at 15,000 ×g for 10 min, and the supernatants were saved for LC-MS/MS analysis.

LC-MS/MS analysis of lipidomics was performed using an Agilent 1290 Infinity HPLC coupled to an Agilent 6540 quadrupole time of flight (QTOF) mass spectrometer. A Waters Acquity CSH C18 column (2.1×100 mm, 1.7 μm) was used for separation. The mobile phase consisted of 60:40 acetonitrile: water (A) and 90:10 isopropanol: acetonitrile (B), both with 10 mM ammonium acetate and 0.1% formic acid. The following gradient was applied: 0-2 min, 15-30% B; 2-2.5 min, 30-48% B; 2.5-11 min, 48-82% B; 11-11.5 min, 82-99% B; 11.5-12 min, 99% B; 12-12.1 min, 99-15% B; 12.1-15 min, 15% B. Column temperature was maintained at 65°C and the flow rate was 0.6 mL/min. MS and MS/MS data were collected in positive and negative ionization mode, with a collision energy of 40 V for MS2. Data processing was performed with MS-DIAL software.

Peak picking, feature alignment and tentative compound annotation were performed with MS-DIAL software. Common lipids were annotated using the LipidBlast *in-silico* MS/MS library in MS-DIAL together with an in-house accurate mass precursor m/z and retention time database. For annotation of peroxidized phospholipid, theoretic m/z was first calculated based on the annotated lipids, with addition of the mass of 2O. Putative annotation was then performed by searching the m/z of unknown peaks against the theoretical m/z, with a mass tolerance of 0.01 Da. Considering their structural similarity, retention times of annotated phospholipids and their peroxidized forms might be close, thus a retention time window of 2 min was applied to exclude potential false positives. Finally, peak heights with lipid annotations were exported and normalized using isotope-labelled internal standards for each lipid class. Theoretical m/z of PE 18:0/20:4 and its peroxidized products were calculated. Peaks were extracted and integrated using Agilent MassHunter software (Ver. B.07.00), with a mass tolerance of 0.01 Da.

### Measurement of hydrogen peroxide production during *in vitro* enzyme assay of POR /CYB5R1

NADPH (200 μM) was incubated with Δ2-42 POR (100 nM) in a Tris-HCl buffer (20 mM Tris, 150 mM NaCl, pH 7.8), after which the reaction buffer was oxygenated (3L/min) for 1 min. After 30 min of incubation, H_2_DCFDA (20 μM) was added into the reaction buffer and the fluorescence changes (λ_ex_= 498/λ_em_=527 nm) were recorded at 20 s intervals approximately 200 times. The quantitative production of hydrogen peroxide by POR was calculated using the fluorescence signal, which corresponds to the fluorescence standard curve of hydrogen peroxide. In some cases, catalase (40,000 units/mg protein, SIGMA) was added to the reaction mixture to deplete the resulting hydrogen peroxide. The curves shown in the figures display one of the three experiments conducted.

Measuring Δ2-28 CYB5R1’s hydrogen peroxide production was performed in a similar manner to Δ2-42 POR, except that NADH was used instead of NADPH.

### Assessment of intracellular H_2_O_2_ levels

A total of 1 x 10^5^ cells per well were seeded on six-well plates one day prior to the experiment. On the next day, cells were treated with different concentrations of H_2_O_2_, then cells were incubated with H_2_DCFDA 500 nM for 1h at 37°C before they were harvested by trypsinization. Subsequently, cells were resuspended in fresh PBS strained through a 70 μM-cut off cell strainer and analyzed using the 488 nm laser of a flow cytometer for excitation. Data was analyzed using FlowJo Software.

### Measurement of superoxide production during *in vitro* enzyme assay of POR

The measurement of superoxide was similar to the process of measuring hydrogen peroxide. The probe hydroethidine (20 μM) was used and the fluorescent signal changes (λ_ex_= 488/λ_em_=567 nm) were recorded at 20 s intervals approximately 200 times.

### The MDA assay to measure lipid peroxidation

A soy phospholipid mixture (100 μg/ml, Avanti) was stored in chloroform, which was full of nitrogen, in a glass bottle at −20 °C. Prior to the lipid peroxidation assay, the phospholipids were dissolved in a 0.05% sodium deoxycholate Tris-HCl buffer (20 mM Tris, 150 mM NaCl, pH 7.8) and sonicated to form a suspension. The phospholipid mixture (1 mg/ml), recombinant Δ2-42 POR (1 μM), and ferric chloride (120 mM) were then added to the Tris-HCl buffer (20 mM Tris, 150mM NaCl, pH 7.8), resulting in final concentrations of 100 µg/ml of phospholipids, 100 nM of Δ2-42 POR, and 120 μM of ferric chloride. The reaction buffer was oxygenated (3L/min) for 1 min, after which 200 μΜ of NADPH was added to the solution to initiate the reaction for 1 h at 37°C. Afterward, 20 µl of the solution was removed from each sample and added to 100 μl of methanol. The mixture was then centrifuged at 15,000 ×g for 5 min and the supernatants were vacuum dried. For the MDA assay, the samples were added to 200 μL of the TBA solution (thiobarbituric acid 240 mg, 7.5 mL of acetic acid, 17.5mL H_2_O) and incubated at 95°C for 60 min. The samples (n=3) were then cooled to room temperature, after which the fluorescence intensity (λ_ex_=532/λ_em_=553 nm) was measured.

The lipid peroxidation assay of Δ2-28 CYB5R1 was similar to that of Δ2-42 POR; the final concentration of Δ2-28 CYB5R1 was 100 nM and NADH replaced NADPH to initiate the reaction.

### Measurement of hydroxyl radicals by DMPO

Recombinant Δ2-42 POR (1 μM), ferric chloride (120 mM), and phospholipids (1 mg/ml) were added to phosphate buffered saline (pH 7.4) to final concentrations of 100 nM Δ2-42 POR; 120 μM ferric chloride; and 100 µg/ml phospholipids. The reaction buffer was oxygenated (3L/min) for 1 min, then NADPH was added to the final concentration of 200 μΜ to initiate the reaction for 1 h at 37°C. Then 1M DMPO (DojinDo) was added to the reaction system to the final concentration of 100 mM. Paramagnetic detection was performed by electron spin resonance paramagnetic spectrometer JEOL-FA200 in Shanghai Institute of Applied Physics, Chinese Academy of Sciences.

### Preparation of liposomes

The soy phospholipid mixture, cardiolipins (Heart, Bovine), 16:0 PC (DPPC), and 18:0-20:4 PE (SAPE) were all obtained from Avanti. The phospholipids were dissolved in chloroform as a 10 mg/ml stock solution at −20°C under nitrogen protection. The phospholipid films were obtained by mixing a 100 μl stock solution and 500 μl of chloroform in round-bottomed 10 ml flasks and evaporating the solvent using a vacuum rotary evaporator (BUCHI). The lipid films were hydrated with 500 μl of buffer L (20 mM HEPES, pH 7.4, 100 mM NaCl) and the liposomes were subjected to 30 rounds of extrusion through a 100 nm polycarbonate membrane using a mini-extruder (Avanti). To prepare the Tb_3+_ encapsulated liposomes, the lipid films were hydrated with 500 μl of buffer TL (20 mM HEPES, pH 7.4, 100 mM NaCl, 50 mM sodium citrate, 15 mM TbCl_3_). After performing the extrusion, the liposomes were washed 8 times with a 100 KD ultrafiltration tube (Millipore) by centrifuging them with buffer L for 20 min at 4°C at a rate of 4,000 × g to remove any external Tb_3+_, after which they were re-suspended in 500 μl of buffer L. All the liposomes were stored at 4°C and used within 24 h.

### Liposome leakage assay

For the duration of the liposome leakage assay, 10 μl of the Tb_3+_ encapsulated liposomes (100 μg/ml) were diluted into 100 μl of buffer L supplemented with 50 μM of DPA. When 100 nM of POR, 50 μM of NADPH, and 120 μM of ferric chloride were added into the reaction mixture in the presence of oxygen, the fluorescence signal (λ_ex_= 270/λ_em_=620 nm) was measured by a microplate reader as Ft_0_. The fluorescence signal was then recorded at 20 s intervals approximately 120 times, after which 0.1% Triton X-100 was added to completely release the Tb_3+_. After adding Triton X-100, the fluorescence signal was measured as Ft_100_. At each time point, the percentage of liposome leakage can be defined as: Leakage (t) (%) = [(Ft-Ft_0_) / (Ft_100_-Ft_0_)] × 100. The curves shown in the figures display one of the three experiments conducted.

### Electron microscopy

POR (100 nM), NADPH (50 μM), and ferric chloride (120 μM) were added to the buffer L, which contained 100 μg/ml of liposome, and was made up of 20% cardiolipin and 80% soy phospholipid mixture. The reaction lasted for 90 mins in the presence of oxygen. Aliquots of the samples (5∼10 μl) were then transferred to carbon support films on electron microscopy grids. After gently washing the unbound liposomes twice with H_2_O, 2% uranyl acetate (5 μl) was negatively stained. The liposomes were imaged on a Tecnai T12 microscope (FEI) at 120 kV and the images were taken on a Gatan 4k × 4k CCD camera with a nominal magnification of 26,000×, resulting in a final pixel size of 4.3 Å.

### Mice xenograft models

Female nu/nu mice aged 4–5 weeks were obtained from Charles River. Luciferase expresing-OVCAR-8 cells were harvested by trypsinization. Subsequently, cells were washed three times with cold PBS and suspended in a 1:1 mixture of PBS and Matrigel (Corning). Each mouse was inoculated subcutaneously with 5 × 10^6^ cells. When tumor volume reached approximately 50 mm^3^, mice were randomly divided into indicated groups. 20mg PACMA31 per kg body weight (10% DMSO, 30% PEG-4000, 60% Saline); 40mg regorafenib per kg body weight (Saline); or 20mg PACMA31 plus 40mg regorafenib per kg body weight daily. PACMA31 was intraperitoneally injected and regorafenib was orally administered. Tumor size was measured using a caliper and weights were determined at the same time. Tumor volume was calculated as follows: (length × width^2^) × 0.5. D-Luciferin (15 mg/mL in PBS, 200μL for each mice) was injected to visualize tumor size *in vivo*. Photographs were captured under IVIS Spectrum.

### Con A-induced acute liver injury mice models

C57BL/6 male mice aged 8–10 weeks were purchased from Vital River Laboratory Animal Technology Co. and housed in a specific pathogen-free animal facility. Mice were injected with 200 μL AAV which expressed shRNA targeting luciferase (shRNA control) or POR (shPOR) via the tail vein. The high titer AAV were packaged by HanBio, China. After 3-4 weeks, the knockdown efficiency in livers of mice were examined by western blotting. For survival studies, the Con A dosage used was 30 mg/kg. In acute liver injury studies, the Con A dosage was reduced to 15 mg/kg. Con A was injected into mice via the tail vein. For measurement of proinflammatory cytokines, blood was collected via mouse eyeball 3 hours after Con A injection. Mice were sacrificed 24 hours post-injection. Livers and blood samples were collected at this time point for H&E staining and measurement of AST/ALT (Abcam), GSH/GSSG (DojinDo), PTGS2 (qPCR, Takara) and MDA (SIGMA) according to the manual’s instructions.

All animal experiments were conducted following the Chinese Ministry of Health national guidelines for the housing and care of laboratory animals and performed in accordance with institutional regulations after review and approval by the Institutional Animal Care and Use Committee of the National Institute of Biological Sciences. The animal experiment was carried out in a blinded manner.

### Synthesis of PACMA31-Probe

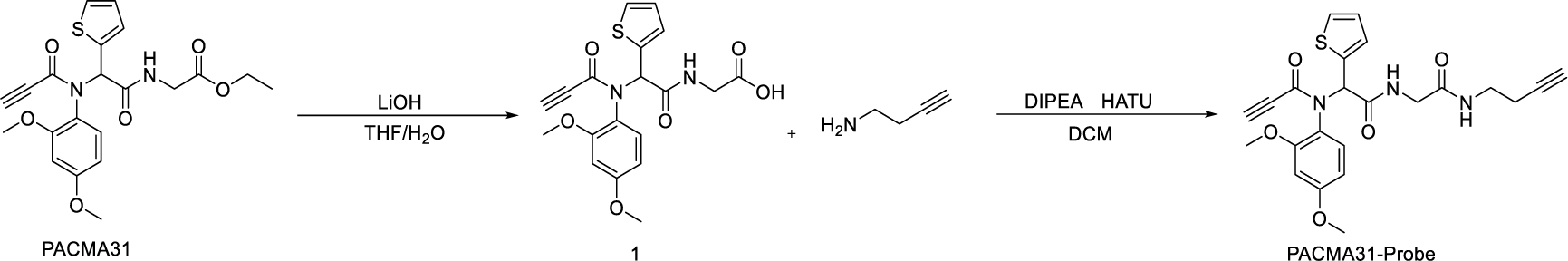

To a solution of PACMA31 (1 g, 2.32 mmol) in the mixture of THF and H_2_O (10 ml, 1:1) LiOH (280 mg, 11.62 mmol) was added and the mixture was stirred at room temperature for 2 h. Then the reaction mixture was poured into water and extracted with dichloromethane (3 x 10 mL) and the organic layer was separated, dried over Na_2_SO_4_, filtered, and concentrated to give compound **1** (900 mg). To a stirred solution of compound **1** (0.5 g, 1.24 mmol) and but-3-yn-1-amine (0.34 g, 4.97 mmol) in dimethylformamide (10 mL) at 0°C, N,N-diisopropylethylamine (0.9 mL, 5 mmol) and HATU (1.9 g, 5 mmol) were added. The mixture was warmed to room temperature and stirred for 24 h. The reaction mixture was put into water (30 mL) and extracted with ethyl acetate (3 x 10 mL). The combined organic layers were washed with saturated brine (20 mL), dried using Na_2_SO_4_, filtered and concentrated *in vacuo*. The residue was purified by chromatography on silica, eluting with methanol (2%)-dichloromethane to give PACMA31-Probe as a white solid (0.32 g, 53%). In CDCl_3_, the mixture of two rotameric forms were in a 6:4 ratio. ^1^H NMR (400 MHz,CDCl3): δ7.65 (t, J=4.0Hz,0.4H), 7.08 (t, J=4Hz,0.6H), 7.23 (d, J=4.0Hz,0.4H), 7.15 (d, J=4.0Hz,0.6H), 7.02 (d, J=4.0 Hz,0.4H), 6.99 (d,J=4.0Hz,0.6H), 6.96 (d, J=4.0Hz,0.6H), 6.91 (d, J=4.0Hz,0.4H), 6.88-6.84 (m, 1×0.6H+1×0.4H), 6.43 (dd, J=8.0Hz,J=4.0Hz, 0.6H), 6.35 (dd, J = 8.0 Hz, J=4.0Hz, 0.4H), 5.79 (s, 0.4H), 5.53 (s, 0.6H), 4.20–4.32 (m, 2×0.6H+2×0.4H), 3.04 (s, 3H), 3.77 (s, 3× 0.4H), 3.78 (s,3×0.6H), 3.40-3.52 (m,2×0.6H+2×0.4H), 2.8 (s, 0.4H), 2.79 (s, 0.6H), 2.412.43 (m, 2× 0.6H+2×0.4H), 1.94–2.00 (m, 1×0.6H+1×0.4H). ^13^CNMR (100 MHz, CDCl3): δ168.2, 161.4, 156.3, 134.5, 131.1, 129.1, 127.1, 126.6, 125.94, 121.46, 104.3, 99.1, 81.5, 79.3, 75.1, 69.6, 62.6, 55.3, 43.0, 38.3, and 19.2.

## Data availability

All of the data support the conclusions relevant to this manuscript are available upon reasonable request from the corresponding authors.

## Acknowledgments

We thank Jingjin Ding, Huan Zeng, Qi Sun, and Yang She from Feng Shao lab for generously sharing with materials for liposome assay and negative staining electron microscopy. We thank Chemistry Center of NIBS for the assistant of drug screen. We thank the Nucleic acid sequencing Center of NIBS for the assistant of gRNA sequencing analysis. We thank Hexia Luo in Electon Microscope Center for assistant of photograph taking. We thank Mr. Alex Wang and Dr. John Hugh Snyder for critical reading of the manuscript. Y.A. was supported by China Postdoctoral Innovation Talent Support Program. This work was supported by institutional grants from the Chinese Ministry of Science and Technology and the Beijing Municipal Commission of Science and Technology. The funders had no roles in study design, data collection, or interpretation of results.

## Author contributions

B.Y., Y.A., and X.W. conceived the study. B.Y. and Y.A. performed most of the experiments. Q.S. helped with the negative staining of liposome for electron microscopy. Y.M. helped with the LC-MS/MS. B.Y., Y.A., Z.Z., and X.W. analyzed and interpreted the results. B.Y., Y.A., and X.W. wrote the manuscript. Z.Z., and X.W. supervised the work.

## Competing interests

The authors declare no competing financial interests.

**Correspondence and requests for materials** should be addressed to B.Y. or Y.A.

## Extended Data Figures

**Extended Data Fig. 1.**
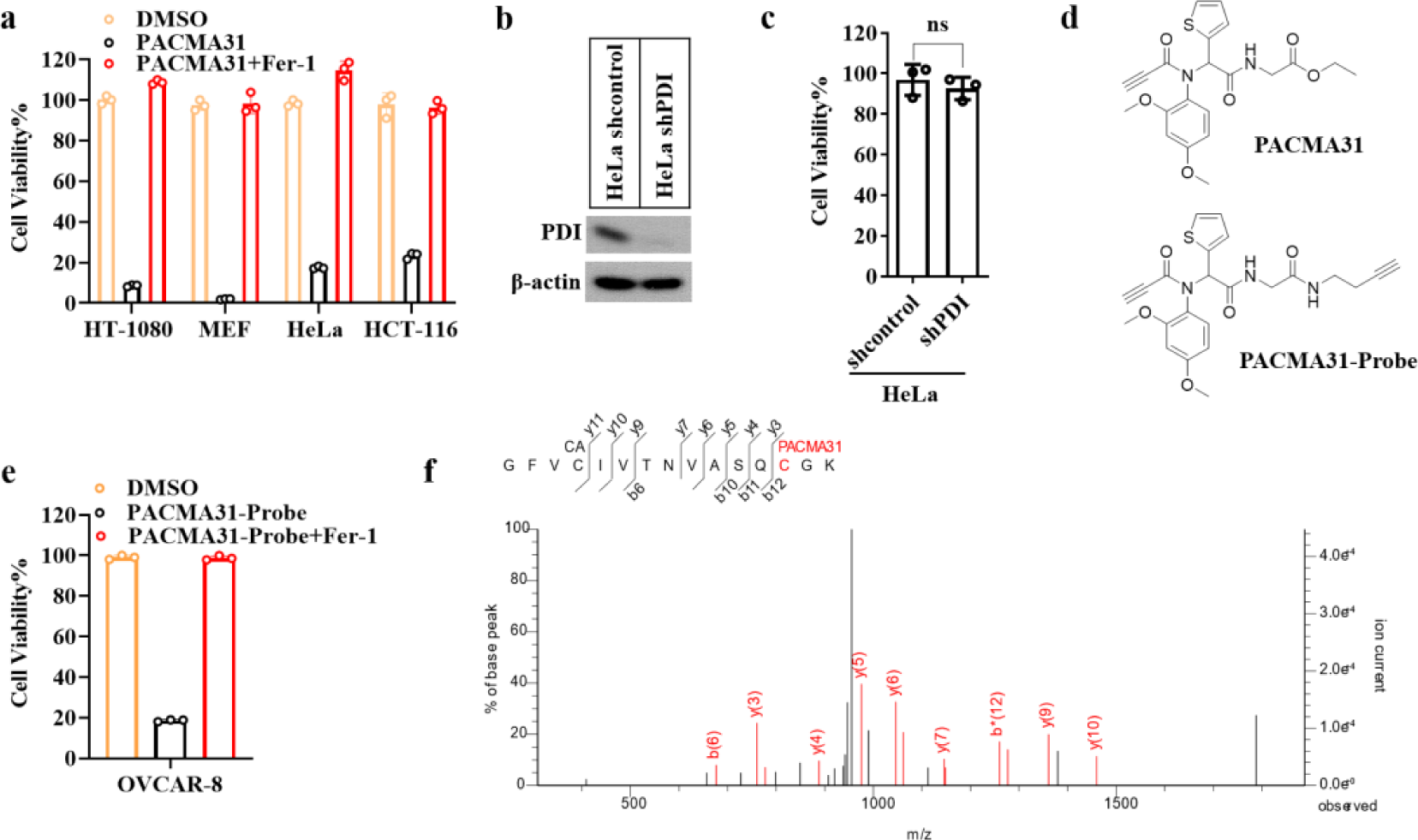
PACMA31 is a novel ferroptosis inducer targeting GPX4. (a) The indicated cell lines were treated with PACMA31 (10 μM) or combined with the ferroptosis inhibitor Fer-1 (1 μM) for 12 hours, followed by monitoring of cell viability. (b) Immunoblot analysis of the PDI protein levels in control or PDI-targeting shRNA treated HeLa cells 36h after the lentiviral infection mediated knockdown. (c) The cell viability of HeLa cells treated with control or PDI-targeting shRNA was examined. (d) The chemical structure of PACMA31 and PACMA31-Probe. (e) The cell viability of OVCAR-8 cells treated with PACMA31-Probe alone or combined with ferroptosis inhibitor Fer-1. (f) The β/γ ion spectra from MS/MS analysis of PACMA31 modified GFVCIVTNVASQCGK peptide establish that PACMA31 covalently modifies Cys-46 of this peptide of GPX4. Cell viability was examined throughout this figure by measuring the ATP level (n=3, repeated in three independent experiments). Student’s *t*-test (two-tailed, unpaired) with mean ± SD: *p < 0.05; **p < 0.01; ***p< 0.001; ns, not significant.

**Extended Data Fig.2.**
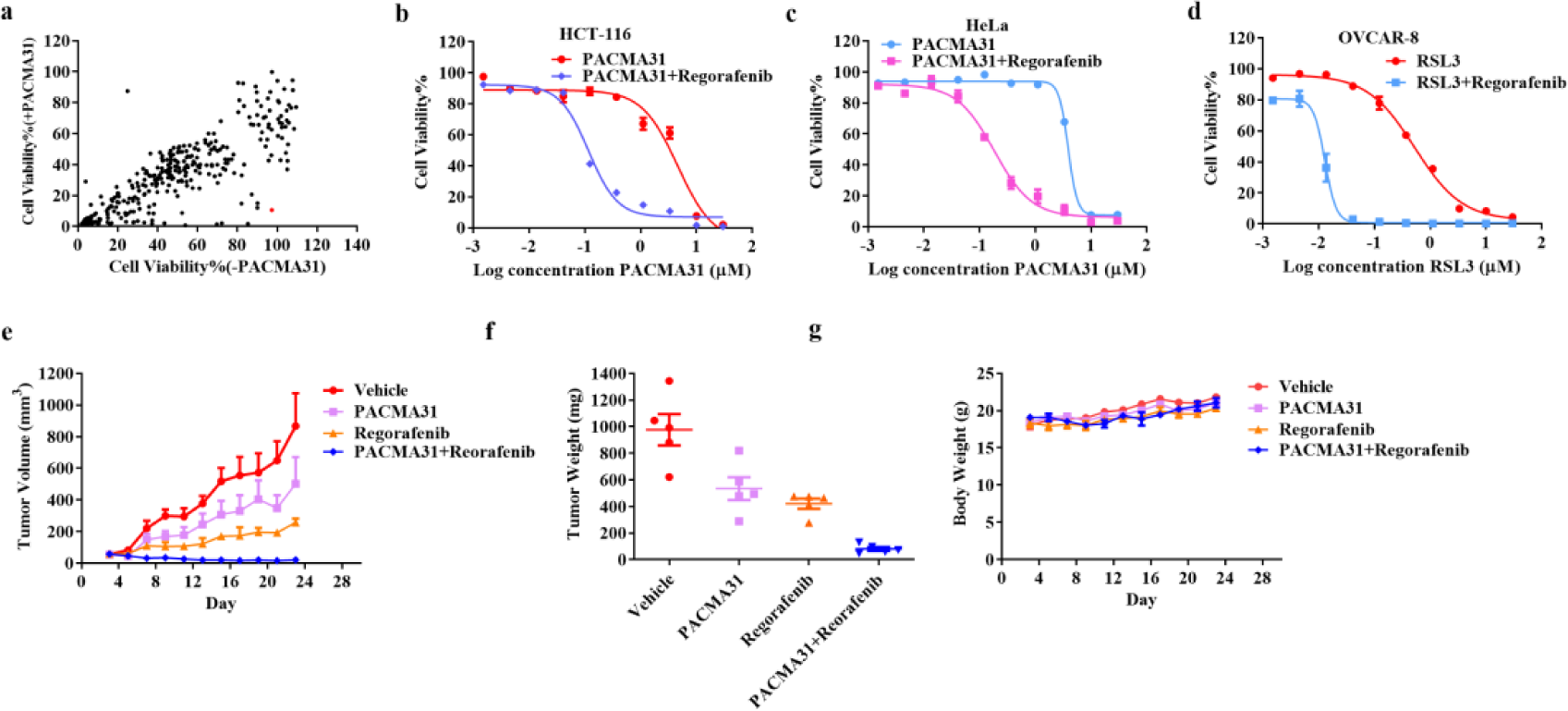
Regorafenib synergistically induces ferroptosis with GPX4 inhibitors. (a) Scatter plot of the cell viability of OVCAR-8 cells treated with different drugs (10μM) in the presence or absence of a subtoxic dose of PACMA31 (200 nM). The red plot represents regorafenib. (b-c) The cell viability curves of HCT-116 (b) or HeLa (c) cells treated with PACMA31 alone or combined with 10 μM regorafenib. (d) The cell viability curves of OVCAR-8 cells treated with RSL3 alone or combined with regorafenib. (e) Tumor growth curves of OVCAR-8 xenograft mice after administration of the indicated drugs for 23 days. (f) Tumor weights of OVCAR-8 xenografts mice after administration of the indicated drugs for 23 days. (g) Body weight curves of OVCAR-8 xenografts mice after administration of the indicated drugs for 23 days. Cell viability assays were examined 12h after treatment by measuring the ATP level using Cell Titer-Glo (n=3, representative results in at least two independent experiments). Student’s *t*-test (two-tailed, unpaired) was used for the comparison of the indicated two groups (mean ± SD).

**Extended Data Fig.3.**
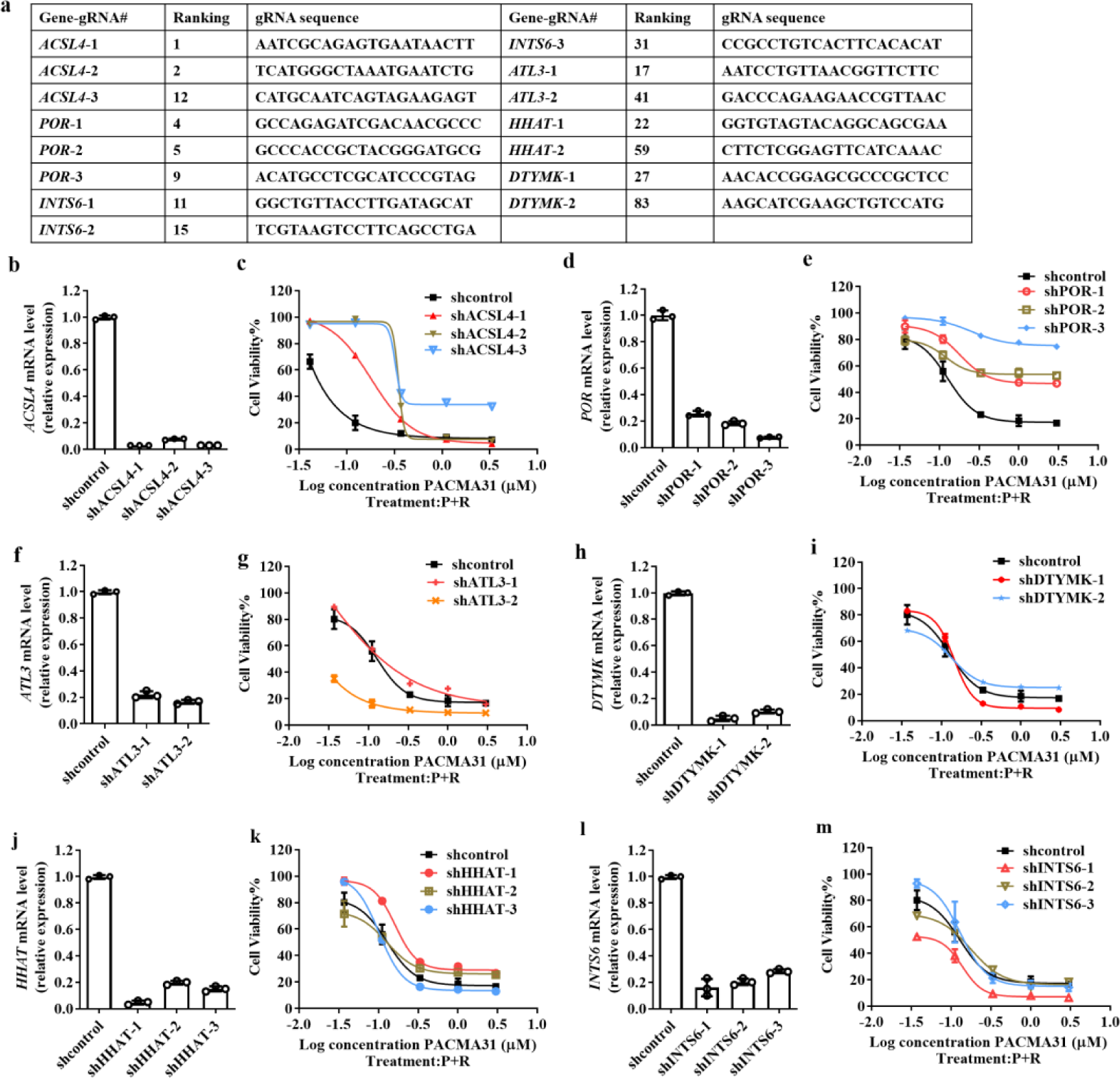
Identification of POR as a novel regulator in P+R induced ferroptosis. (a) The gRNA hits from a genome-wide CRISPR-Cas9 screen of P+R-induced ferroptosis in HeLa-Cas9 cells. Fold-enrichment-based ranking and the target sequences of each gRNA were listed. The genes with at least 2 gRNAs involved in the top 100 gRNAs were selected as candidates. (b) The relative mRNA levels of ACSL4 in HeLa cells treated with control or shRNA targeting ACSL4. (c) The cell viability of HeLa cells treated with control or shRNA targeting ACSL4 followed by P+R. (d) The relative mRNA levels of POR in HeLa cells treated with control or shRNA targeting POR. (e) The cell viability of HeLa cells treated with control or shRNA targeting POR followed by P+R. (f) The relative mRNA levels of HeLa cells treated with control or shRNA targeting ATL3. (g) The cell viability of HeLa cells treated with control or shRNA targeting ATL3 followed by P+R. (h) The relative mRNA levels of DTYMK in HeLa cells treated with control or shRNA targeting DTYMK. (i) The cell viability of HeLa cells treated with control or shRNA targeting DTYMK followed by P+R. (j) The relative mRNA levels of HHAT in HeLa cells treated with control or shRNA targeting HHAT. (k) The cell viability of HeLa cells treated with control or shRNA targeting HHAT followed by P+R. (l) The relative mRNA levels of INTS6 in HeLa cells treated with control or shRNA targeting INTS6. (m) The cell viability of HeLa cells treated with control or shRNA targeting INTS6 followed by P+R. Cell viability was repeated three times by measuring the ATP levels using Cell Titer-Glo (n=3). All bar graphs show the mean ± SD.

**Extended Data Fig.4.**
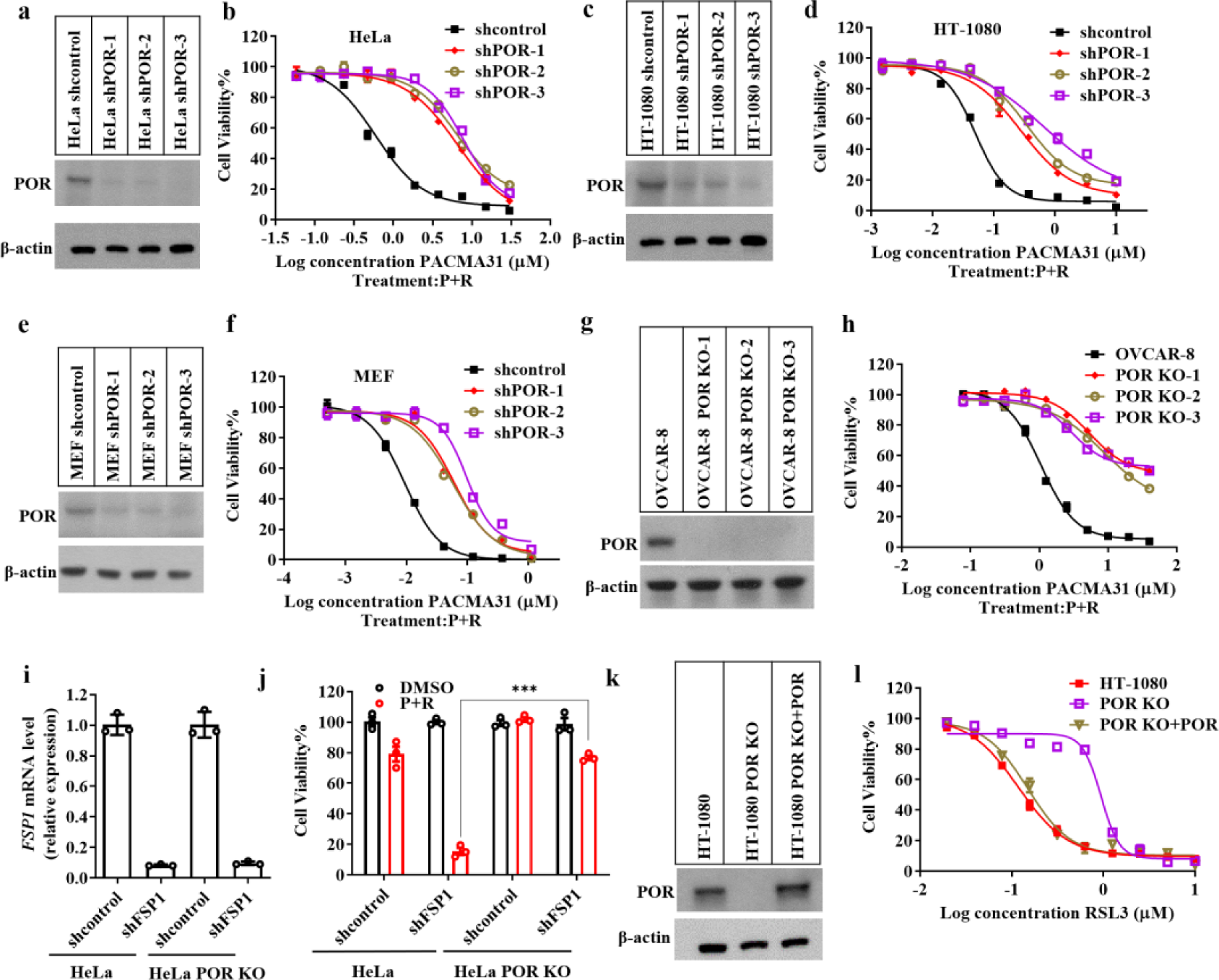
POR is a general regulator in ferroptosis. (a) Immunoblot analysis of the POR protein levels in HeLa cells treated with control or shRNA targeting POR. (b) The cell viability curves in HeLa cells treated with control or shRNA targeting POR followed by P+R. (c) Immunoblot analysis of the POR protein levels in HT-1080 cells treated with control or shRNA targeting POR. (d) The cell viability curves in HT-1080 cells treated with control or shRNA targeting POR followed by P+R. (e) Immunoblot analysis of the POR protein levels in MEF cells treated with control or shRNA targeting POR. (f) The cell viability curves of MEF cells treated with control or shRNA targeting POR followed by P+R. (g) Immunoblot analysis of the POR protein levels in OVCAR-8 cells (parental or POR KO). (h) The cell viability curves in OVCAR-8 cells (parental or POR KO) treated with P+R. (i) The relative mRNA levels of FSP1 in HeLa cells (parental or POR KO) treated with control and shRNA targeting FSP1 to determine knockdown efficiency using quantitative PCR (n=3). (j) The cell viability of HeLa cells (parental or POR KO) treated with control and shRNA targeting FSP1 followed by P+R (PACMA31 200 nM; Regorafenib 10 μM). (k) Immunoblot analysis of the POR protein levels in HT-1080 (parental, POR KO, POR KO stably expressed with POR) cells. (l) The cell viability curves of HT-1080 (parental, POR KO, POR KO stably expressed with POR) cells in response to treatment of P+R. Cell viability was repeated three times with three wells each time. Student’s *t*-test (two-tailed, unpaired) with mean ± SD: *p < 0.05; **p < 0.01; ***p< 0.001; ns, not significant.

**Extended Data Fig.5.**
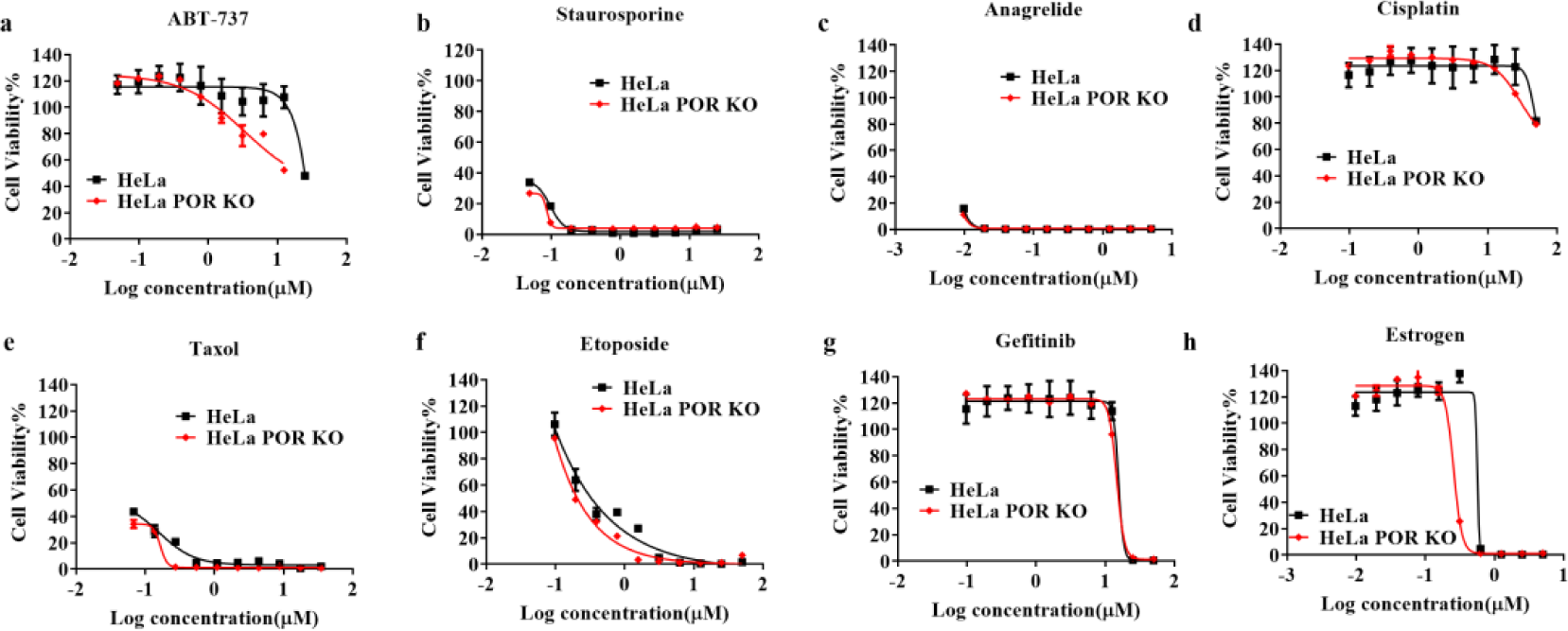
POR is not involved in selected cytotoxic compounds induced cell death. The cell viability of HeLa (parental or POR KO) cells treated with Bcl-2/Bcl-X_L_ dual inhibitor ABT-737 (a), apoptosis-inducing pan-kinase inhibitor staurosporine (b), apoptosis-inducing, PDE3-inhibitor anagrelide (c), cisplatin (d), Taxol (e), etoposide (f), gefitinib (g), estrogen (h). Cell viability was measured in this figure by measuring the ATP level using Cell Titer-Glo (n=3, repeated for three times). All bar graphs throughout the figure show the mean ± SD.

**Extended Data Fig.6.**
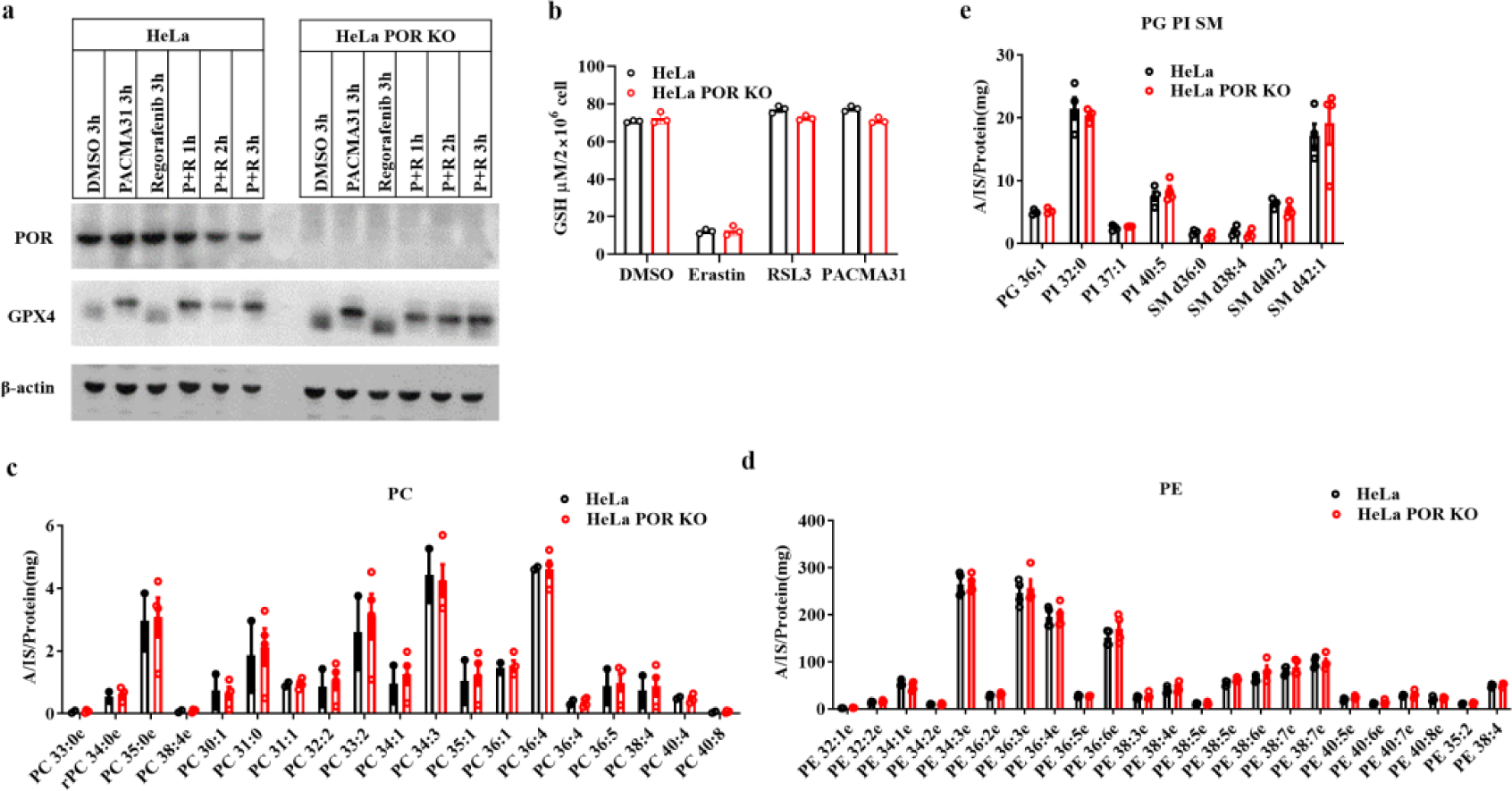
POR knockout does not change GPX4 expression, GSH levels, and phospholipids composition of HeLa cells. (a) Immunoblot analysis of the GPX4 protein levels in HeLa (parental or POR KO) cells with indicated treatments (PACMA31 1 μM; regorafenib 10 μM). (b) The GSH levels of HeLa (parental or POR KO) cells with indicated treatments. Erastin (10 μM) was incubated with cells for 12h, PACMA31 (1 μM) or RSL3 (1 μM) was incubated for 1h. (c-e) Lipidomic profile of HeLa (parental or POR KO) cells. The data represents the mean value of the ratio of the area of analyte (A)/internal standard (IS)/protein (mg). Each group has three replicates, and results were representative of two independent experiments.

**Extended Data Fig.7.**
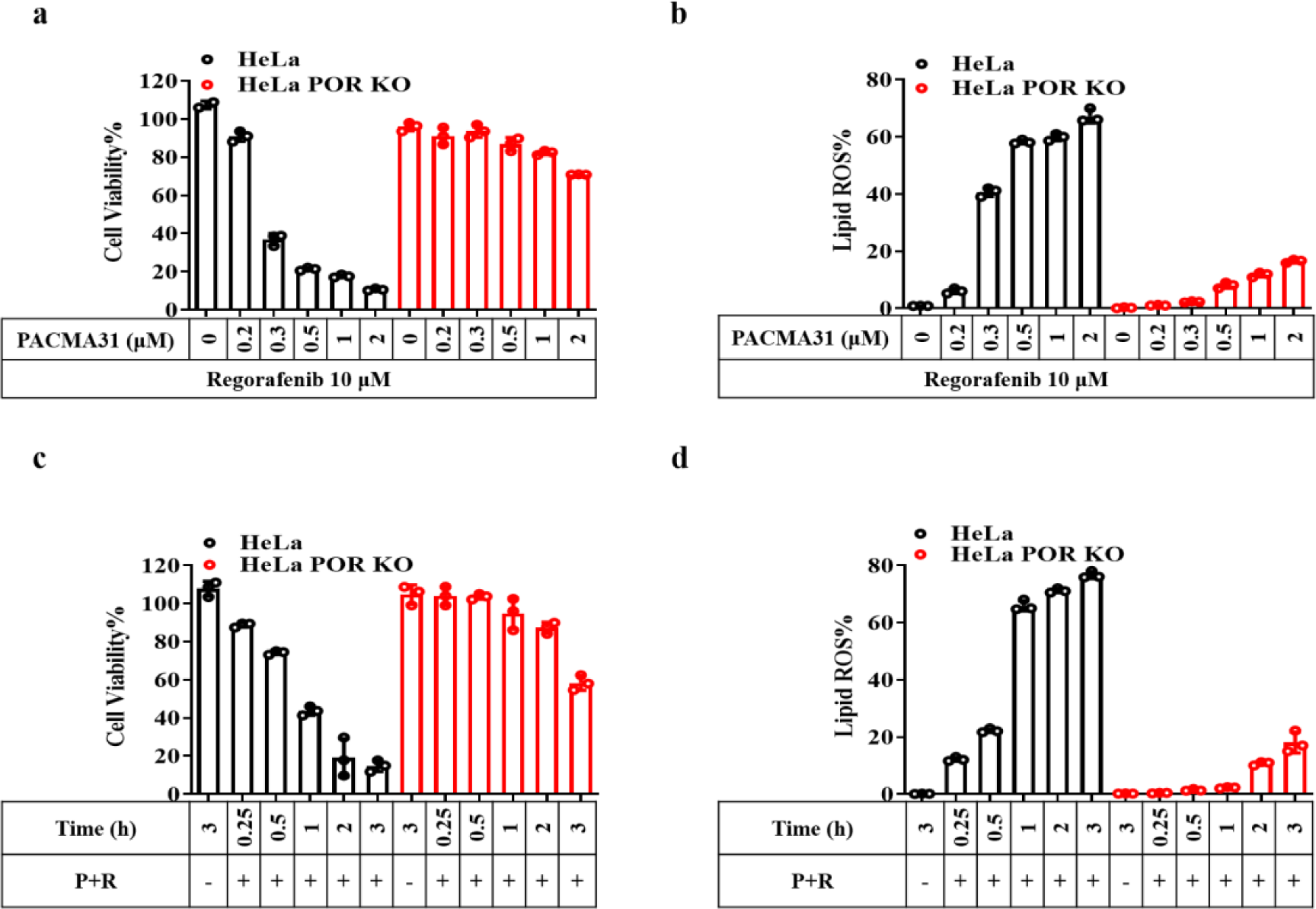
POR was required for lipid peroxidation during ferroptosis. (a-b) HeLa (parental or POR KO) cells were treated with different does of PACMA31 combined with regorafenib (10μM) for 3h. The cell viability was determined by measuring ATP levels (a). Lipid peroxidation was measured by Liperfluo and analyzed by flow cytometry (b). (c-d) HeLa cells (parental or POR KO) were treated with P+R (PACMA31 2 μM; Regorafenib 10 μM) for the indicated time points. The cell viability was determined (c), and lipid peroxidation was detected based on the oxidation of the probe Liperfluo, assessed via flow cytometry (d). Bar graphs show the mean ± SD. Experiments were repeated three times with three replicates each time.

**Extended Data Fig.8.**
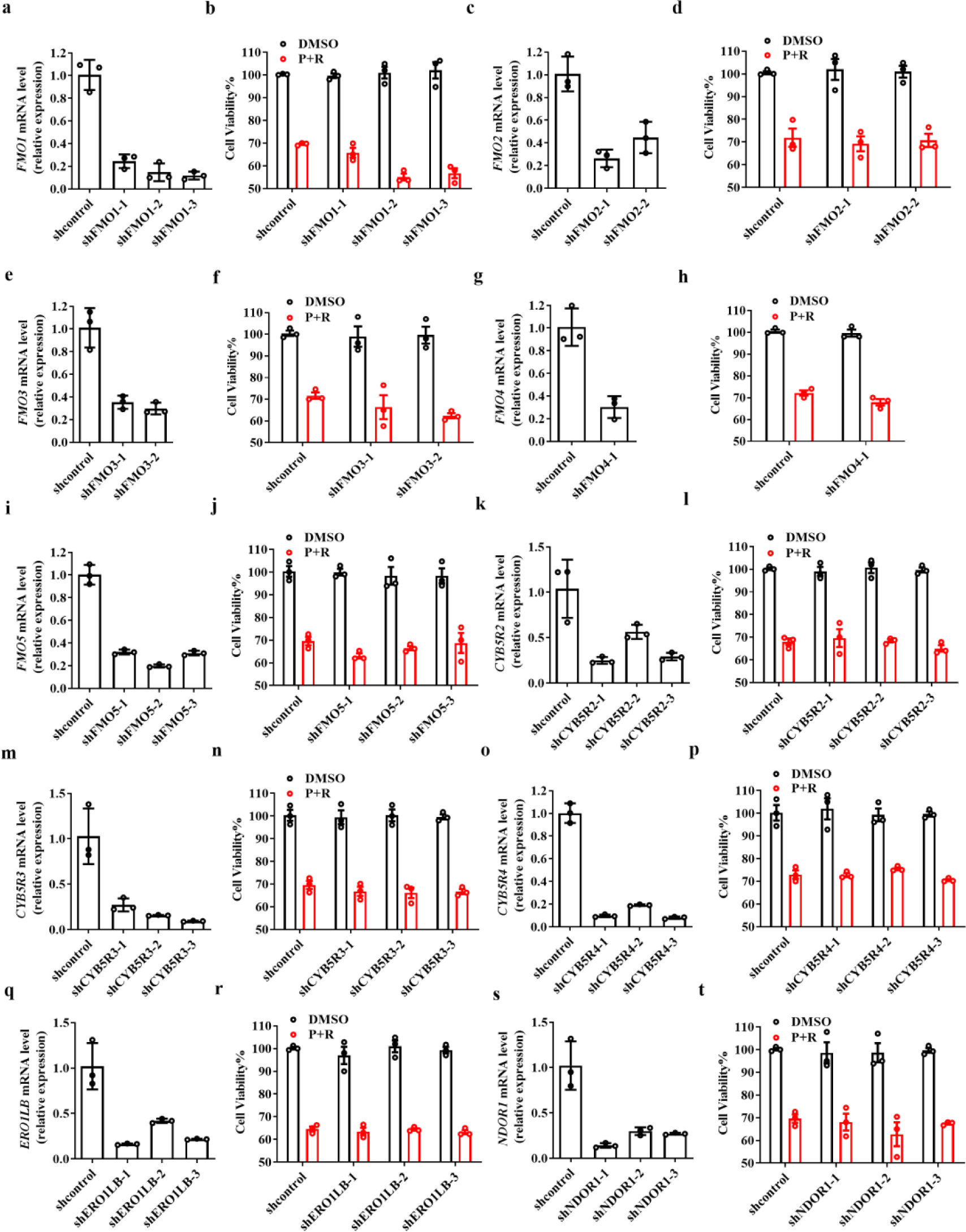
Other oxidoreductases do not involve in ferroptosis. The reductases including: FMO1 (a-b); FMO2 (c-d); FMO3 (e-f); FMO4 (g-h); FMO5 (i-j); CYB5R2 (k-l); CYB5R3 (m-n); CYB5R4 (o-p); ERO1LB (q-r); NDOR1 (s-t) were knocked down with shRNA species specifically targeting mRNAs of these proteins. The mRNA levels of these enzymes were determined by qPCR (n=3). Cell viability throughout the figure was examined after 12h by measuring the ATP level using Cell Titer-Glo. (PACMA31 2 μM; regorafenib 10 μM; n=3; repeated for two times). Bar graphs show mean ± SD.

**Extended Data Fig.9.**
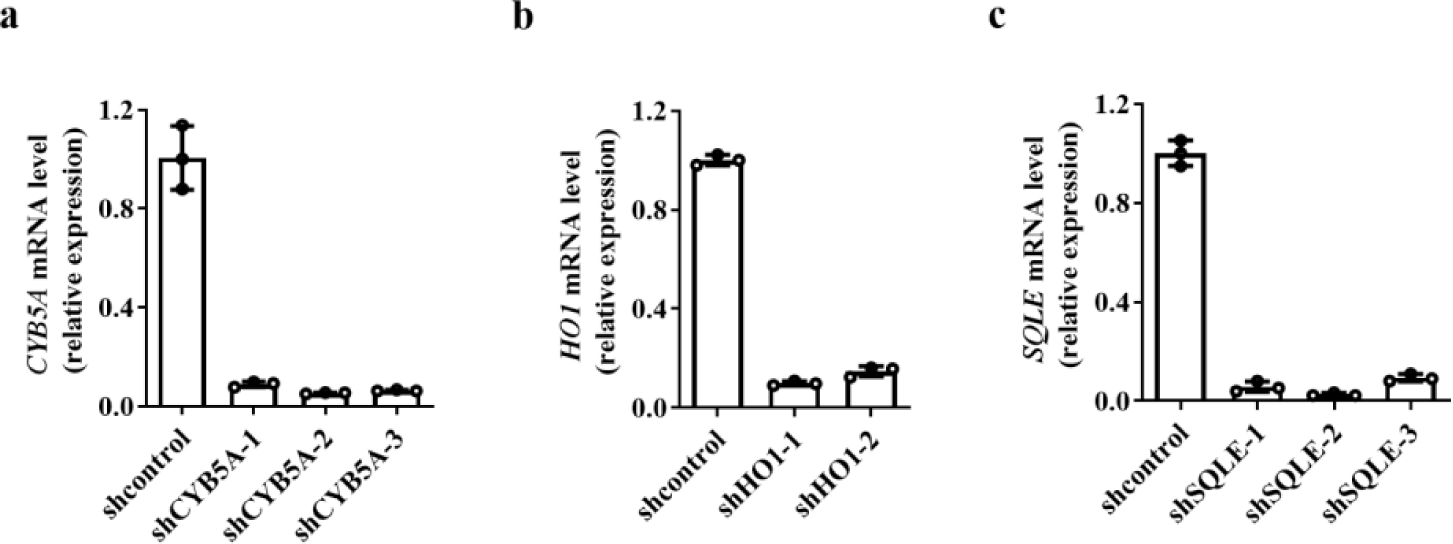
Interaction with downstream partner proteins is dispensable for POR and CYB5R1’s function in ferroptosis. (a) The relative mRNA levels of CYB5A in HeLa treated with control or shRNA targeting CYB5R1. (b) The relative mRNA levels of SQLE in HeLa (shRNA control or shSQLEs) cells. (c) The relative mRNA levels of HO1 in HeLa (shRNA control or shHO1s) cells. Results were repeated two times, with three replicates each time. Student’s *t*-test (two-tailed, unpaired) with mean ± SD: *p < 0.05; **p < 0.01; ***p< 0.001; ns, not significant.

**Extended Data Fig.10.**
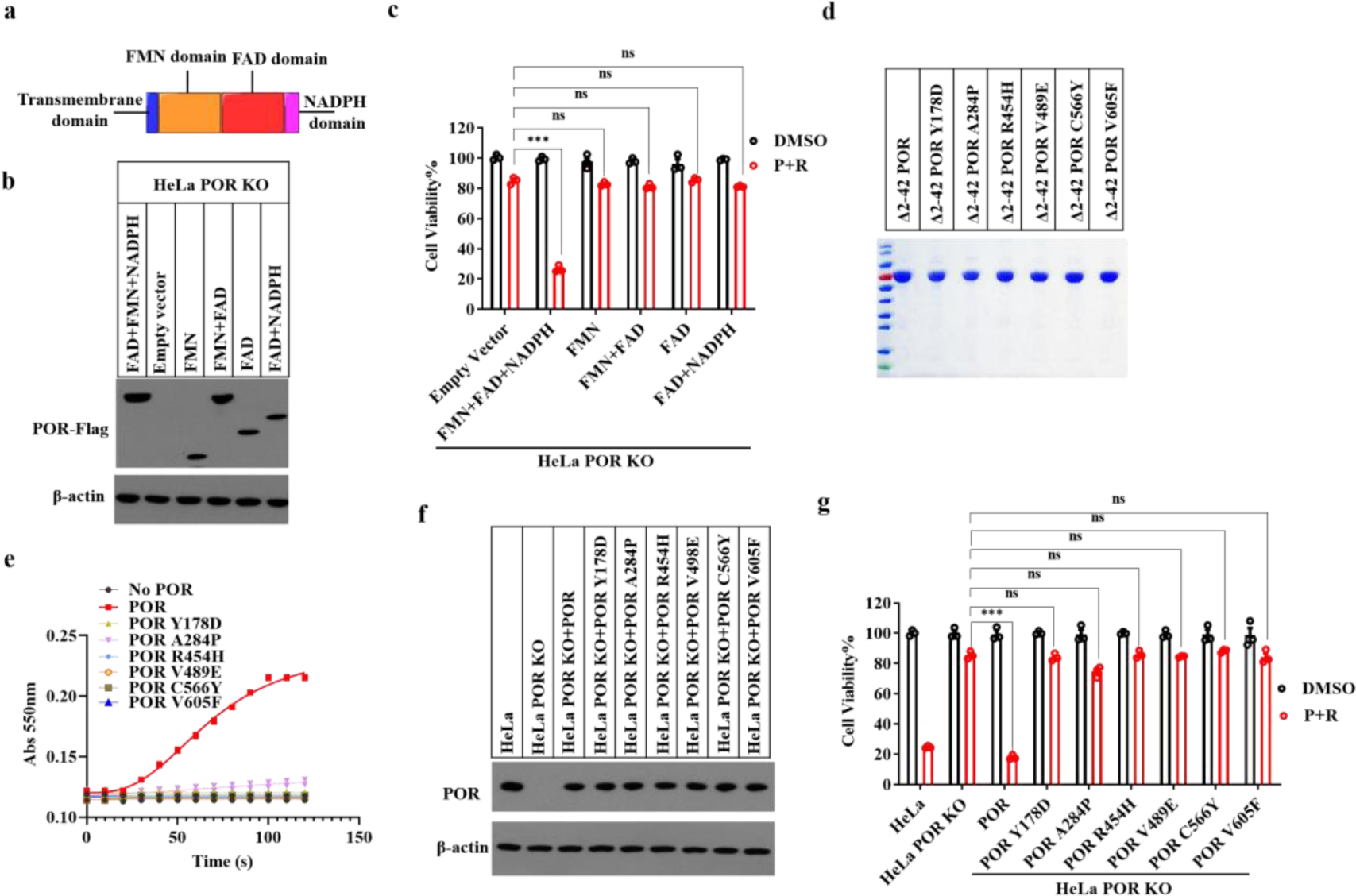
The electron transfer ability was required for POR’s function in ferroptosis. (a) Schematic diagram of the various functional domains of POR. (b) Immunoblot analysis of different truncations of POR expressed in HeLa POR KO cells. The Flag-tag was introduced into C-terminal of all the truncations. (c) The cell viability of HeLa POR KO cells expressing different POR truncations under P+R (PACMA31 1 μM; Regorafenib 10 μM) induced ferroptosis. (d) SDS-PAGE and Coomassie brilliant blue stain of recombinant Δ2-42 POR and different site mutants of POR proteins. (e) The reduction of cytochrome c in the present of POR or indicated variants of POR. Reduced cytochrome c were detected in 550 nm (absorbance). (f) Immunoblot analysis of the POR protein levels in HeLa POR KO cells expressing POR and indicated POR variants. (g) The cell viability of HeLa POR KO cells expressing POR and indicated POR variants treated with P+R (PACMA31 1 μM; Regorafenib 10 μM). Cell viability was examined after 12h by measuring the ATP level (n=3; repeated for three times). Student’s *t*-test (two-tailed, unpaired) was used for the comparison of the indicated two groups (mean ± SD): *p < 0.05; **p < 0.01; ***p< 0.001; ns, not significant.

**Extended Data Fig.11.**
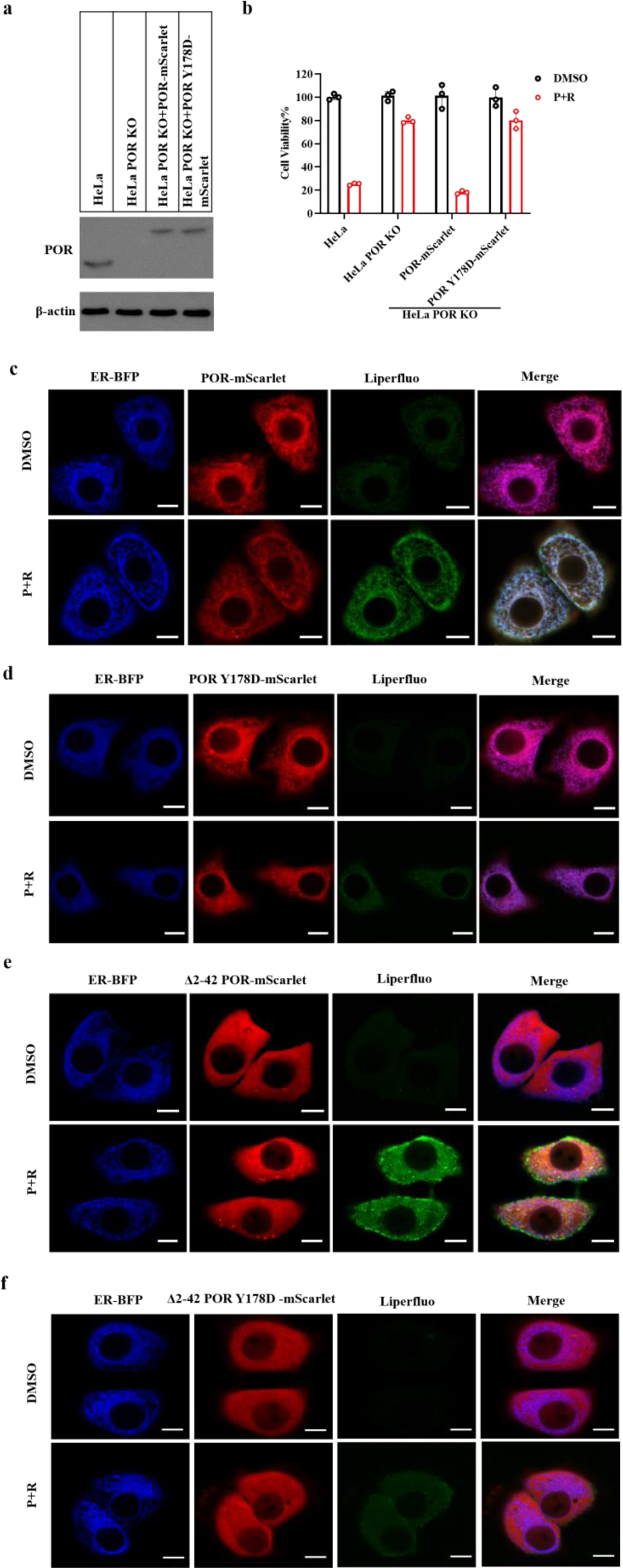
The electron transfer activity of POR is required for lipid peroxidation. (a) Immunoblot analysis of the POR protein in parental and HeLa POR KO cells stable integrated with empty vector or POR-mScarlet variants vectors (The mScarlet was introduced into C-terminal of POR). (b) The cell viability of parental or HeLa POR KO cells stable integrated with empty vector or POR-mScarlet variants vectors treated with P+R (PACMA31 1 μM; regorafenib 10 μM) for 12h. (c-d) Fluorescence confocal microscopy of HeLa POR KO cells expressing ER-BFP-KDEL (ER localized) and POR-mScarlet (c) or POR Y178D-mScarlet (d) treated with P+R for 2h. The lipid peroxidation was detected by probe Liperfluo. (e-f) The fluorescence confocal microscopy of HeLa POR KO cells expressing ER-BFP-KDEL and Δ2-42 POR-mScarlet (e) or Δ2-42 POR Y178D-mScarlet (f) treated with or without P+R for 2h. The lipid peroxidation was detected by probe Liperfluo. Scale bars, 10 μm. Cell viability throughout the figure was examined after 12h by measuring the ATP level with mean ± SD (n=3). All the data were representative of three independent experiments.

**Extended Data Fig.12.**
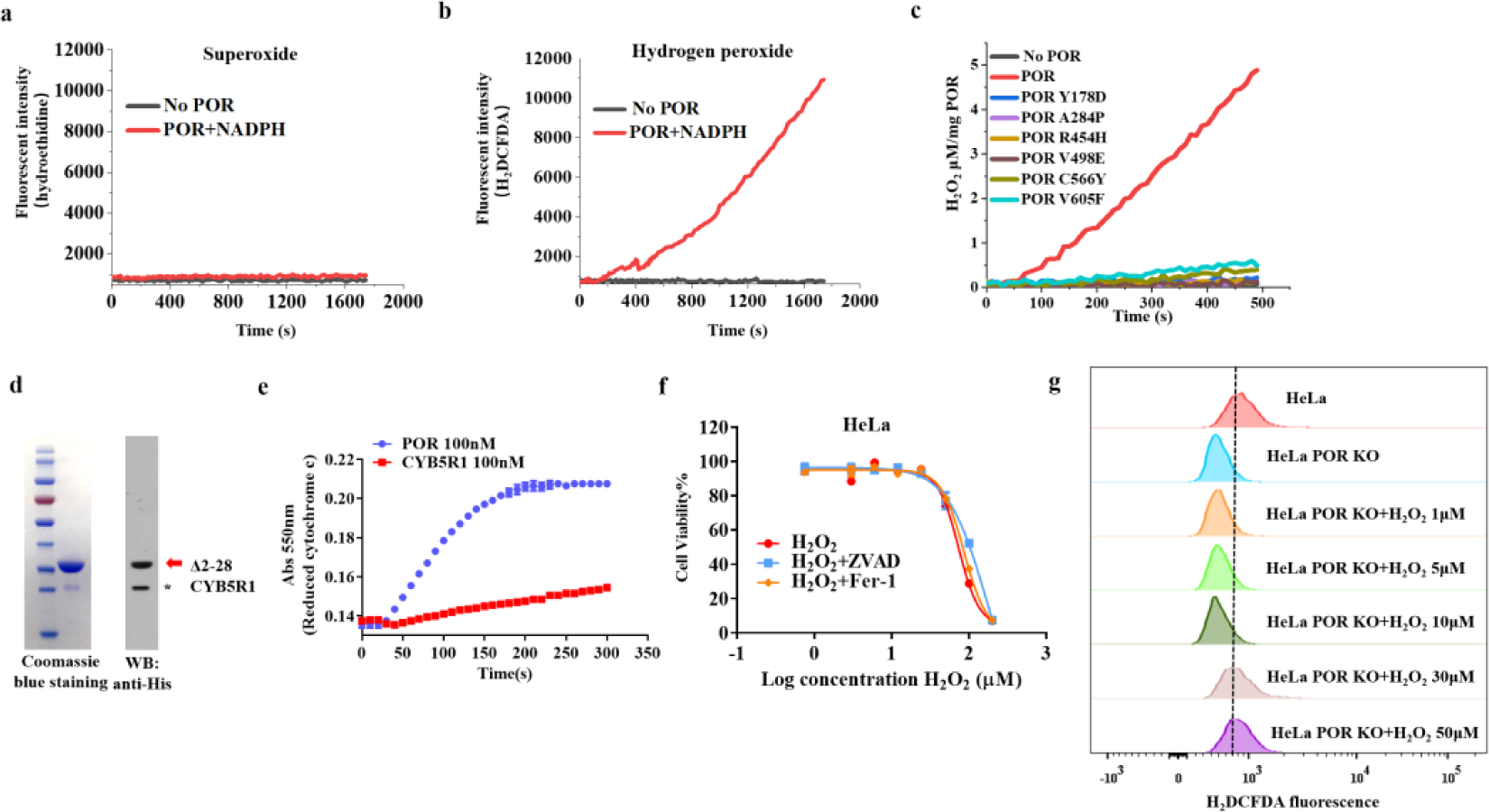
POR catalyzes the production of hydrogen peroxide. (a) Fluorescence changes of the reaction buffer containing POR and NADPH after sequential additions of hydroethidine (for detection of superoxide). (b) Fluorescence changes of the reaction buffer containing POR and NADPH after sequential additions of H_2_DCFDA (for detection of hydrogen peroxides). (c) The hydrogen peroxide production rates of different POR variants. (d) SDS-PAGE with Coomassie brilliant blue staining and western blotting (against His tag) of recombinant Δ2-28 CYB5R1 (red arrow) purified from bacteria. Asterisk indicates a His-CYB5R1 fragment. (e) The reduction of cytochrome c in the present of POR or CYB5R1. (f) Cell viability curves of HeLa cells treated with different concentrations of H_2_O_2_ in the absence or presence of the caspase inhibitor ZVAD or the ferroptosis inhibitor Fer-1. (g) The intracellular H_2_O_2_ level of HeLa and HeLa POR KO cells (treated with the indicated concentrations of H_2_O_2_ in culture medium) were analyzed by flow cytometry. Cell viability throughout the figure was examined after 12h by measuring the ATP level with mean ± SD (n=3). All the data were representative of three independent experiments.

**Extended Data Fig.13.**
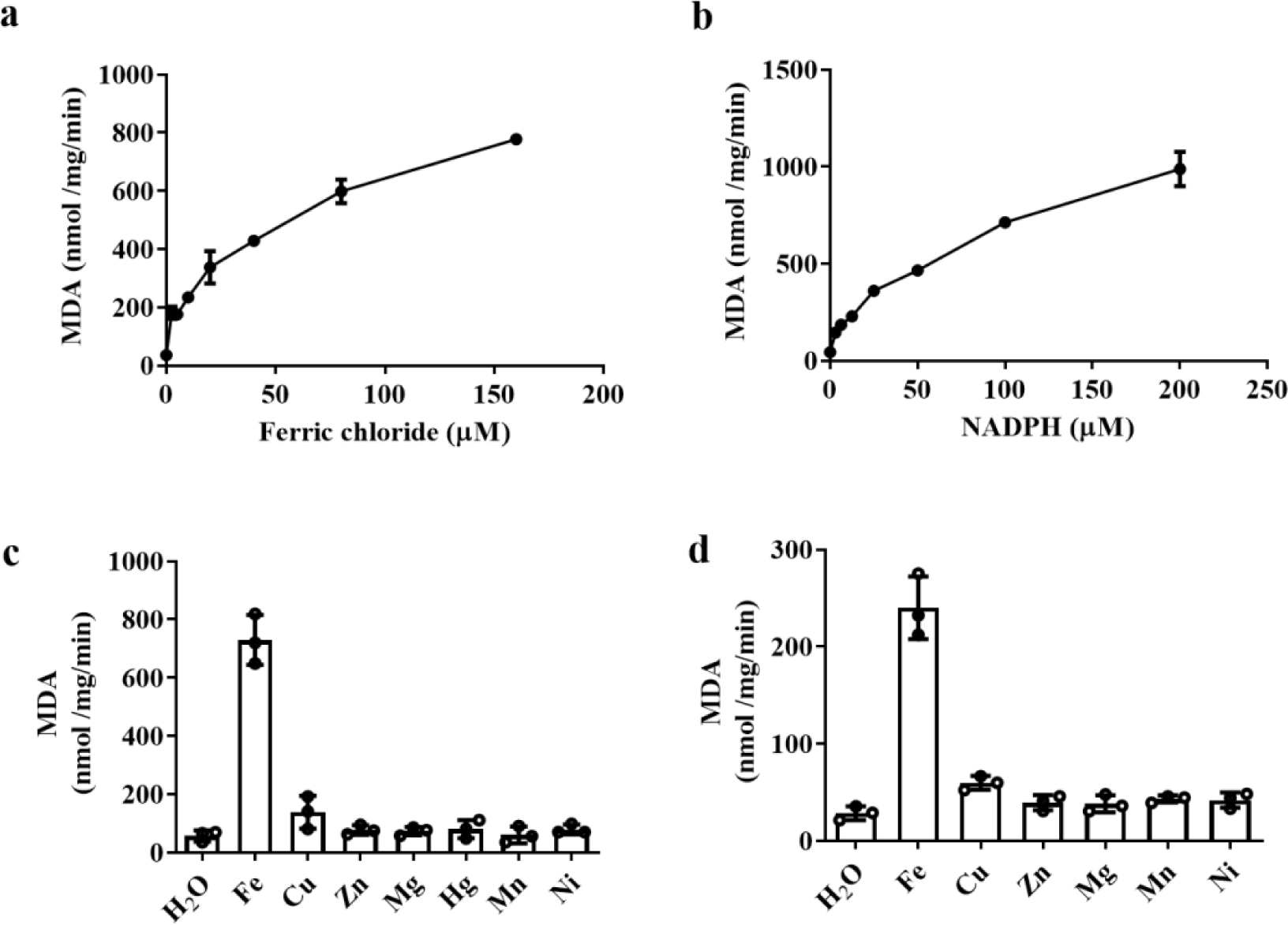
POR and CYB5R1 catalyze lipid peroxidation. (a) The production of MDA in the presence of POR (100 nM) with increasing concentrations of ferric chloride. Other reaction components included 200 μM NADPH and 100 μg/ml phospholipids mixture. (b) The production of MDA in the presence of POR with increasing concentrations of NADPH. (c) The production of MDA in the presence of POR with different metal ions (Ferric chloride, Zinc sulfate monohydrate, Copper (II) sulfate pentahydrate, Mercury (II) sulfate, Nickel (II) sulfate hexahydrate, Magnesium sulfate heptahydrate, Manganese (II) chloride, 120 μM each). (d) The production of MDA in the presence of CYB5R1 with indicated metal ions (120 μM each). The data were representative results of three independent experiments. Bar graphs show mean ± SD.

**Extended Data Fig.14.**
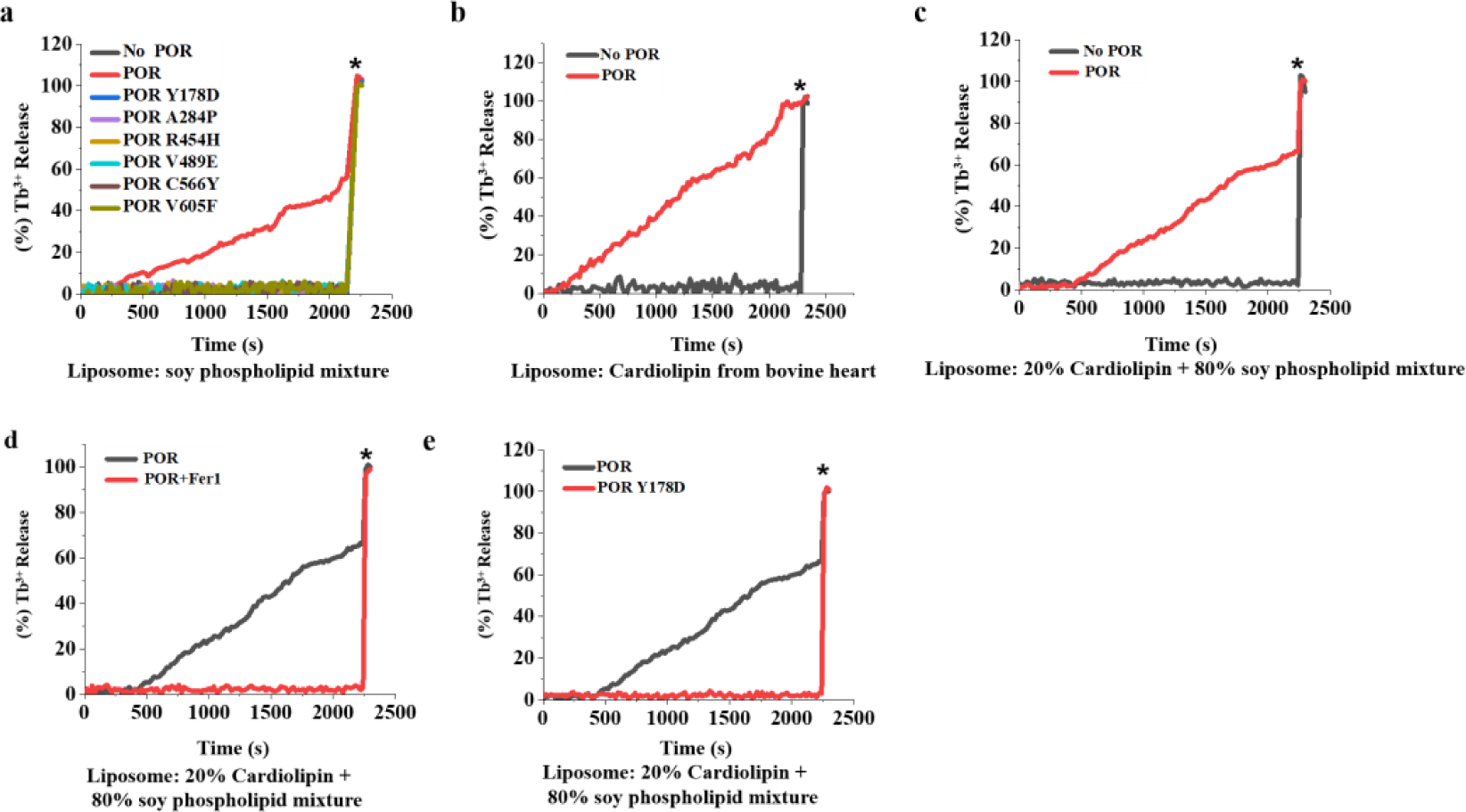
POR and CYB5R1 disrupt phospholipids membrane and cause liposome leakage. (a) Time courses of liposome leakage induced by different POR variants (100 nM). The liposome was made by soy phospholipids mixture. Asterisks indicated the time points when Triton X-100 was added to achieve complete release of Tb_3+_. (b) Time courses of liposome leakage in the present of POR. Liposomes constitute from bovine heart cardiolipins. (c) The liposome leakage in the presence of POR. Liposomes constitute of 20% cardiolipin from bovine heart and 80% phospholipids mixture from soybean. (d) The liposome leakage induced by POR in the absence or presence of Fer-1 (10 μM). (e) The liposome leakage induced by POR or POR Y178D mutant. Results were representative of three independent experiments. Experiments in figures c-e were performed in one plate and shared the same controls.

**Extended Data Fig.15.**
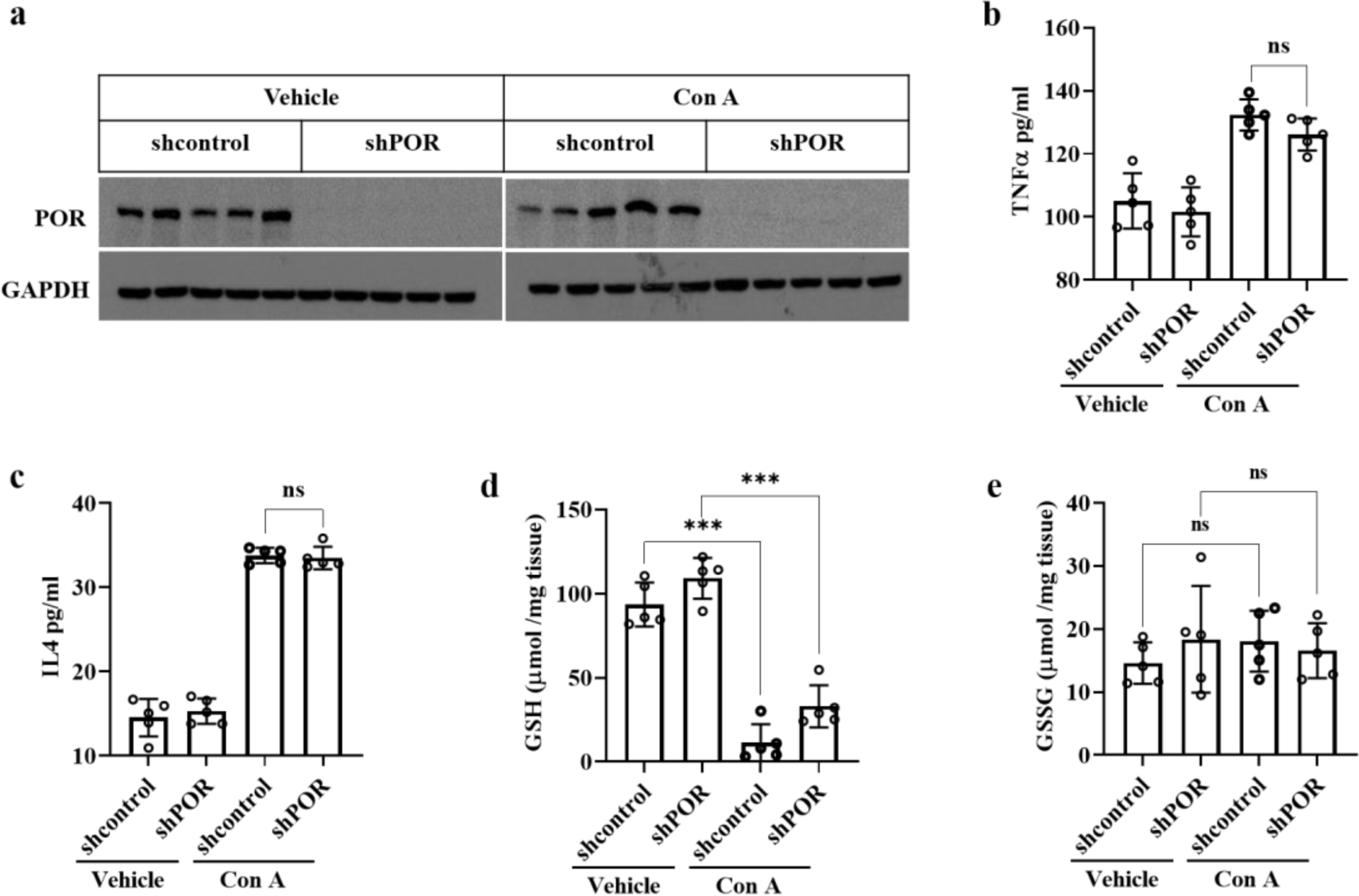
POR knockdown had no effect on the increase of proinflammatory cytokines and decrease of GSH levels in Con A-induced liver injury mice models. (a) POR expression level in the liver of shRNA control and shPOR mice. (b) Serum levels of TNFα in mice measured by ELISA. (c) Serum levels of IL4 in mice measured by ELISA. (d) Liver GSH levels of indicated groups. (e) Liver GSSG levels of indicated groups.

**Extended Data Fig.16.**
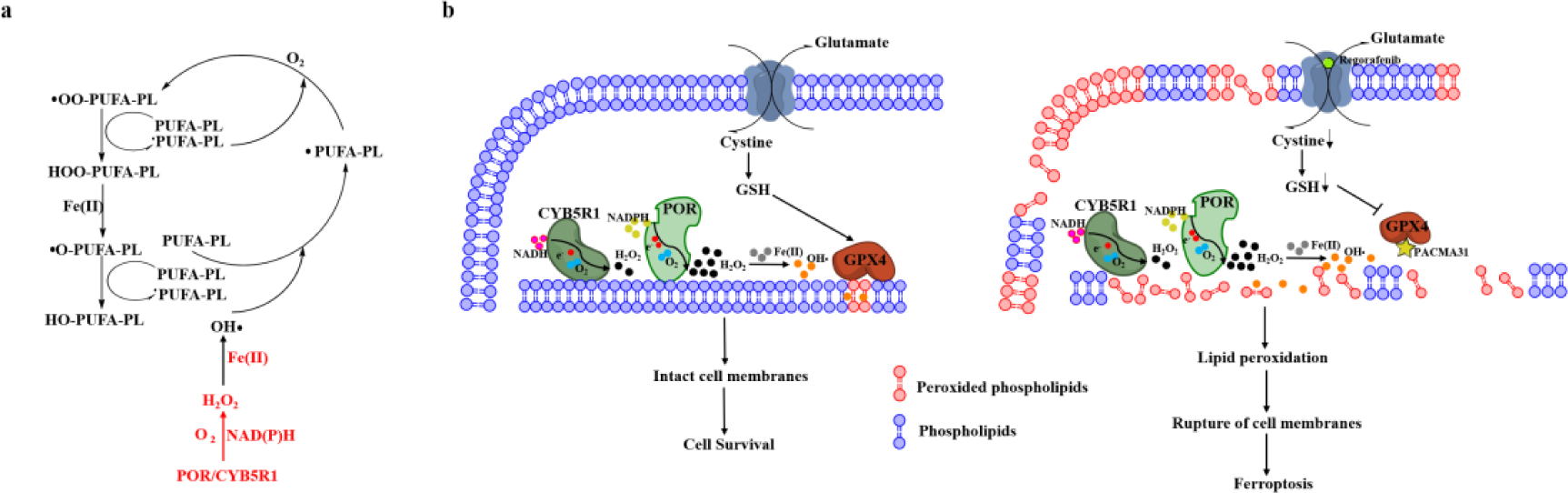
A schematic model describing the roles of POR/CYB5R1 in regulating lipid peroxidation and ferroptosis. (a) Scheme of the reaction mechanism of POR/CYB5R1 catalyzed lipid peroxidation. The lipid peroxidation begins with the hydrogen peroxide produced by POR/CYB5R1. Hydrogen peroxide reacts with ferrous ion to produce hydroxyl radicals, which can abstract the bis-allylic hydrogen, rearrange the resonance radical structure in PUFA phospholipids. Then oxygen was added into the radicals leading to the formation of peroxyl radicals, and then the hydroperoxyl phospholipids are formed. (b) In normal conditions, POR/CYB5R1 can catalyze lipid peroxidation as a consequence of its electron transfer process, but the produced lipid peroxides are typically eliminated by GPX4/FSP1 antioxidant systems to maintain cell survival. Upon disruption of GPX4 antioxidant systems, the lipid peroxides produced by POR/CYB5R1 accumulate, eventually levels high enough to initiate ferroptosis.

